# No evidence of impaired sensorimotor adaptation in Complex Regional Pain Syndrome

**DOI:** 10.1101/2020.09.08.287862

**Authors:** Axel D. Vittersø, Gavin Buckingham, Antonia F. Ten Brink, Monika Halicka, Michael J. Proulx, Janet H. Bultitude

**Author notes:** Corresponding author, Address: Department of Psychology, 10 West, University of Bath, Claverton Down, Bath, BA2 7AY, United Kingdom, URL: https://www.bath.ac.uk/research-centres/centre-for-pain-research-cpr/. **CrRediT author statement: Axel Vittersø:** Conceptualization, Methodology, Software, Formal analysis, Investigation, Resources, Data Curation, Writing - Original Draft, Visualization, Project administration, Funding acquisition. **Gavin Buckingham:** Methodology, Software, Writing - Review & Editing, Supervision, Funding acquisition. **Antonia Ten Brink:** Investigation, Resources, Writing - Review & Editing. **Monika Halicka:** Resources, Writing - Review & Editing. **Michael Proulx:** Writing - Review & Editing, Supervision, Funding acquisition. **Janet Bultitude:** Conceptualization, Methodology, Resources, Writing - Review & Editing, Supervision, Funding acquisition.

## Abstract

Sensorimotor conflict is theorised to contribute to the maintenance of some pathological pain conditions, such as Complex Regional Pain Syndrome (CRPS). We therefore tested whether sensorimotor adaptation is impaired in people with CRPS by characterising their adaption to lateral prismatic shifts in vision. People with unilateral upper limb CRPS Type I (n = 17), and pain-free individuals (n = 18; matched for age, sex, and handedness) completed prism adaptation with their affected/non-dominant and non-affected/dominant arm, in a counterbalanced order. We examined 1) the rate at which participants compensated for the optical shift during prism exposure (i.e. strategic recalibration), 2) endpoint errors made directly after prism adaptation (sensorimotor realignment) and their retention, and 3) kinematic markers associated with feedforward motor control and sensorimotor realignment. We found no evidence that strategic recalibration was different between people with CRPS and controls, including no evidence for differences in a kinematic marker associated with trial-by-trial changes in movement plans. Participants made significant endpoint errors in the prism adaptation after-effect phase, which are indicative of sensorimotor realignment. Overall, the magnitude of this realignment was not found to differ between people with CRPS and pain-free controls. However, people with CRPS made greater endpoint errors when using their affected hand than their non-affected hand, whereas no such difference was seen in controls. Taken together, these findings suggest that strategic control and sensorimotor realignment were not impaired for either arm in people with CRPS. In contrast, they provide some evidence that there is a greater propensity for sensorimotor realignment in CRPS, consistent with more flexible representations of the body and peripersonal space. Our study challenges the theory that sensorimotor conflict might underlie pathological pain that is maintained in the absence of tissue pathology.

## 1. Introduction

Complex Regional Pain Syndrome (CRPS) is characterised by pain, motor deficits, and autonomic symptoms (Harden et al., 2010; Harden, Bruehl, Stanton-Hicks, & Wilson, 2007). In addition to its physical manifestation, CRPS can be accompanied by a range of neuropsychological changes (for reviews, see Halicka, Vittersø, Proulx, & Bultitude, 2020a; Kuttikat et al., 2016). These changes include distorted representations of the body (e.g. Lewis, Kersten, McCabe, McPherson, & Blake, 2007; Moseley, 2005; Peltz, Seifert, Lanz, Müller, & Maihöfner, 2011), and altered updating of such representations (Vittersø, Buckingham, Halicka, Proulx, & Bultitude, 2020). Sensorimotor processing might also be affected by altered sensory experiences and motor deficits (Harden et al., 2010; Harden et al., 2007).

Harris (1999) proposed that incongruence between motor predictions and sensory outcomes, such as might arise in CRPS, could underlie pathological pain conditions for which there is no clear tissue damage (see also McCabe & Blake, 2007; McCabe, Blake, & Skevington, 2000). Experimental manipulations that induce sensorimotor conflict have been found to increase pain and anomalous sensations in people with CRPS (Brun et al., 2019) and fibromyalgia (Brun et al., 2019; Martínez, Guillen, Buesa, & Azkue, 2019; McCabe, Cohen, & Blake, 2007), whereas the evidence is mixed for whiplash associated disorders (Daenen et al., 2012; Don, De Kooning, et al., 2017). Such manipulations can also increase sensory anomalies without altering pain levels in people with arthritis (Brun et al., 2019), low back pain (Don et al., 2019), dancers with musculoskeletal pain (Roussel et al., 2015), or for violinists with pain (Daenen, Roussel, Cras, & Nijs, 2010; for review, see Don, Voogt, Meeus, De Kooning, & Nijs, 2017). For people with such conditions, problematic levels of sensorimotor incongruence could arise in daily life from compromised motor predictions and altered sensory feedback. Under normal circumstances, however, adaptation occurs when faced with conflicting information that appears to originate from the same source (e.g. in time and space; Wei & Kording, 2009). That is, the sensorimotor system can detect the discrepancy between actual and intended outcome of a movement, and adjust the spatial mappings of sensory inputs and/or the motor command for future movements to reduce the conflict (Wolpert, Diedrichsen, & Flanagan, 2011). If sensorimotor conflict underlies CRPS and related conditions, then it follows that this adaptation process could be disrupted such that the sensorimotor system is unable to compensate for incongruent information.

Prism adaptation is a useful paradigm for investigating sensorimotor adaptation because it enables scrutiny of several distinct sensorimotor processes (e.g. strategic recalibration, sensorimotor realignment, and retention; Fig. 1). These processes have been studied in great detail (for review see Jacquin-Courtois et al., 2013) and are associated with distinct cortical mechanisms (for review see Panico, Rossetti, & Trojano, 2020). A typical prism adaptation procedure involves performing ballistic pointing movements while wearing goggles fitted with prismatic lenses that create a lateral optical shift (Held & Freedman, 1963; Redding, Rossetti, & Wallace, 2005; Von Helmholtz, 1924). During prism exposure, participants initially make pointing errors in the direction of the prismatic shift. These pointing errors will quickly reduce as movements are repeated, such that pointing is once again accurate within about a dozen trials. At first, the pointing errors are reduced mainly through “strategic recalibration”, a somewhat conscious process in which participants deliberately adjust their aim or mentally rotate the target location to correct for the visual shift (Rossetti, Koga, & Mano, 1993). In the longer term (e.g. over 50-100 movements) the spatial reference frames that coordinate visual, motor, and proprioceptive processing gradually realign to compensate for the optical distortion introduced by the prisms) (“sensorimotor realignment” ; Jeannerod & Rossetti, 1993; Redding et al., 2005). Once the prism goggles are removed, people will typically make pointing errors in the direction opposite to the optical displacement (the prism adaptation “after-effect”). With further repeated pointing, these errors will quickly reduce if they are visible to the participants (e.g. the hand is not occluded). Even after an extended washout period, a small degree of the prism adaptation after-effects can still be observed when pointing movements are performed without visual feedback (retention). The retention of prism adaptation after-effects reflects the degree to which the realignment of visual and proprioceptive reference frames is maintained (Prablanc et al., 2019).

**Figure 1.**
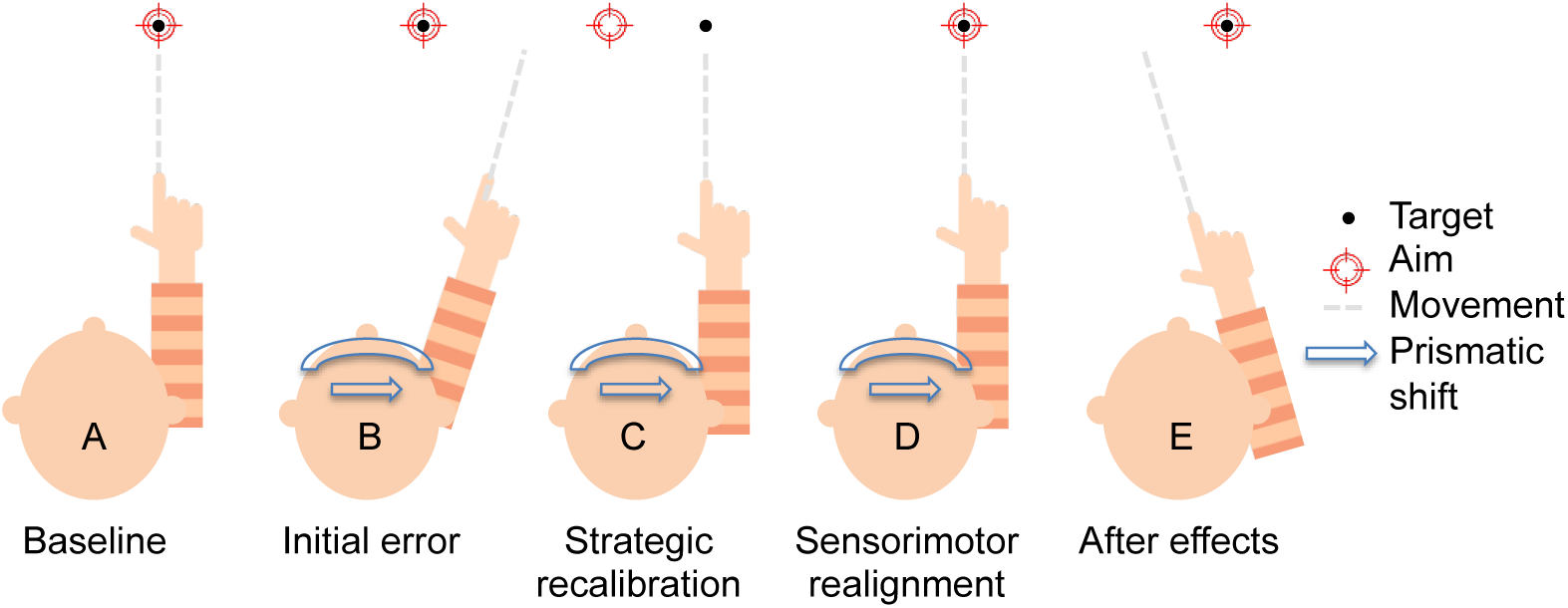
The processes involved in prism adaptation are depicted. Baseline pointing errors (A) and prism after-effects (E) are measured without wearing goggles. During early prism exposure trials (B) participants make initial errors in the direction of the prismatic shift (to the right in this example). In the first few trials of prism exposure participants correct their pointing mainly through strategic recalibration (e.g. deliberately aiming to the left of target; C). Over the longer term, sensorimotor realignment occurs (D) such that the represented visual location of the target is brought into alignment with the felt location of the arm, and participants no longer need to deliberately mis-aim to reach the target. Once the goggles are removed participants make errors in the direction opposite to the prismatic shift (i.e. after effects; E), which are leftward in this example.

The distinct sensorimotor processes involved in prism adaptation can be quantified by measuring endpoint errors and kinematic markers, and can be expressed with exponential decay functions. Strategic recalibration (Fig. 1C) can be measured by the reduction in endpoint errors during early prism exposure. Sensorimotor realignment (Fig. 1D) and its retention are typically indexed by the magnitude of the endpoint errors made once the goggles are removed (i.e. during open-loop pointing; Fig. 1E), relative to baseline (Prablanc et al., 2019). In order to quantify the rapid changes during early prism exposure, endpoint errors can be fitted to an exponential decay function (Facchin, Bultitude, Mornati, Peverelli, & Daini, 2018; Martin, Keating, Goodkin, Bastian, & Thach, 1996a; Nemanich & Earhart, 2015; O’Shea et al., 2014). Some degree of online correction is common during prism exposure, which can mask some of the rapid changes that occur. One way to circumvent this problem is to examine the pointing movement prior to any visual feedback being available to the participant. People will typically update their aim to compensate for the error made on a previous trial, which can be observed in the direction in which they initiate a movement (O’Shea et al., 2014). Therefore, kinematic recordings of arm movements during prism exposure allow for the trial-by-trial changes in movement plans to be computed, independently of any online control, which shed light on the process of strategic recalibration. Kinematic recordings also allow endpoint errors to be accurately measured, and are therefore advantageous when studying the distinct sensorimotor processes involved in prism adaptation.

We aimed to investigate sensorimotor adaptation in pathological pain by characterising prism adaptation in people with upper-limb CRPS compared to pain-free controls. We investigated strategic recalibration, and the development, magnitude, and retention of sensorimotor realignment. Participants underwent prism adaptation once with each hand, which enabled us to compare outcomes between Groups (CRPS or controls), and the Hand used (affected/non-dominant, non-affected/dominant). We hypothesised that people with CRPS would show impaired strategic recalibration (Hypothesis 1) and sensorimotor adaptation (Hypothesis 2). That is, we expected that compared to pain-free controls, people with CRPS would require more trials for endpoint errors to asymptote during prism exposure, and would show smaller magnitudes and less retention of after-effects. We also hypothesised that the sensorimotor realignment might develop and/or decline at a different rate for people with CRPS, compared to controls (Hypothesis 3). We hypothesized that people with CRPS would show less evidence of trial-by-trial changes in movement plans to compensate for the prismatic shift than controls (Hypothesis 4). The trial-by-trial changes allow for deliberate changes in movement plans to be measured, which are an important strategy involved in strategic calibration (O’Shea et al., 2014), whilst eliminating any contribution from online corrections. As certain deficits (e.g. proprioceptive; Bank, Peper, Marinus, Beek, & van Hilten, 2013) can be apparent in the healthy limb, although more subtle than the CRPS-affected limb, we also considered that any deficits in strategic control, sensorimotor realignment, and/or trial-by-trial changes in movement would be limited to, or more apparent in, the CRPS-affected arm compared to the non-affected arm (Hypothesis 5).

## 2. Materials and methods

We used a single-session mixed design in which participants with CRPS and controls each underwent prism adaptation using each arm in the same session, and we compared the performance between the two arms and between groups. The study was preregistered on the Open Science Framework (https://osf.io/6jpfg/).

### 2.1. Participants

Seventeen people with CRPS type 1 predominantly affecting one upper limb (*M*_age_ = 53.53 years, *SD* = 11.67; 16 female; 14 right-handed; 9 left-affected CRPS; Table 1) were recruited through the UK national CRPS registry and from our own database. The latter is an internal database of people with CRPS consenting to be contacted about research who have been referred to us from the Royal United Hospitals (Bath, UK) and Oxford University Hospitals (Oxford, UK), and Royal National Orthopaedic Hospital (London, UK) NHS Foundation Trusts; or who have contacted us directly. We decided on our sample size pragmatically, based on the maximum number of people with CRPS we could feasibly recruit and test given financial and time constraints. Twelve participants met the Budapest research criteria for CRPS (Harden et al., 2010; Harden et al., 2007), three met the clinical criteria, and two were diagnosed with CRPS not otherwise specified. Fourteen of the people with CRPS had previously participated in a randomised control trial of prism adaptation for pain relief in which half of the participants underwent twice daily prism adaptation treatment and half performed identical movements while wearing goggles fitted with neutral lenses (Halicka, Vittersø, et al., 2020b; Halicka, Vittersø, Proulx, & Bultitude, 2020b). There was an average of 15.13 months (*SD* = 6.97) between participants completing the exposure phase of the randomised control trial and when they took part in the current study. The remaining participants with CRPS had never undergone prism adaptation before. Eighteen pain-free control participants (*M*_age_ = 54.17 years, *SD* = 12.22; 17 female; 15 right-handed) who were matched to the participants with CRPS for age (±5 years), sex, and self-reported handedness were recruited from a community sample. None of the pain-free control participants had undergone prism adaptation before.

**Table 1.**
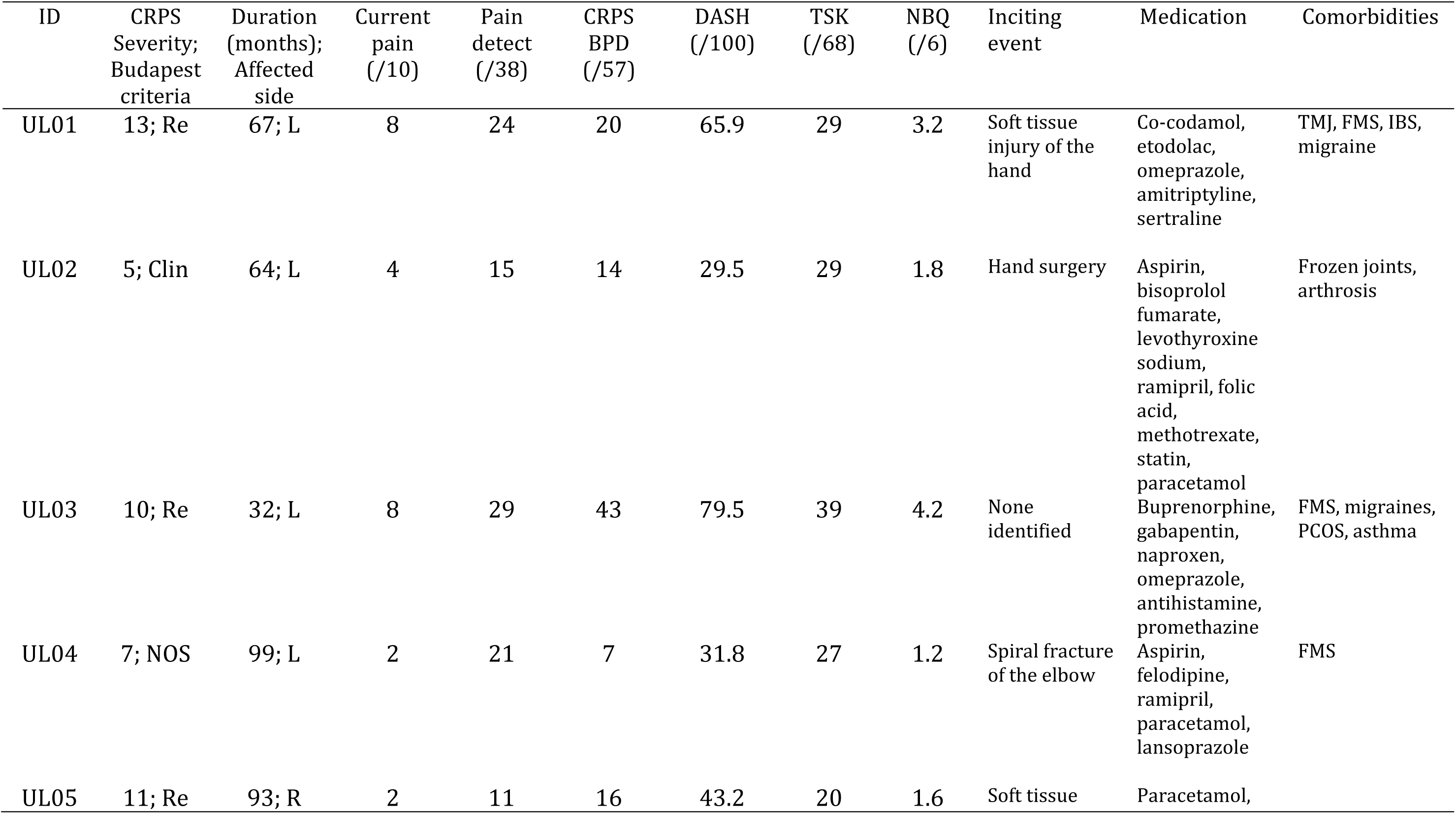
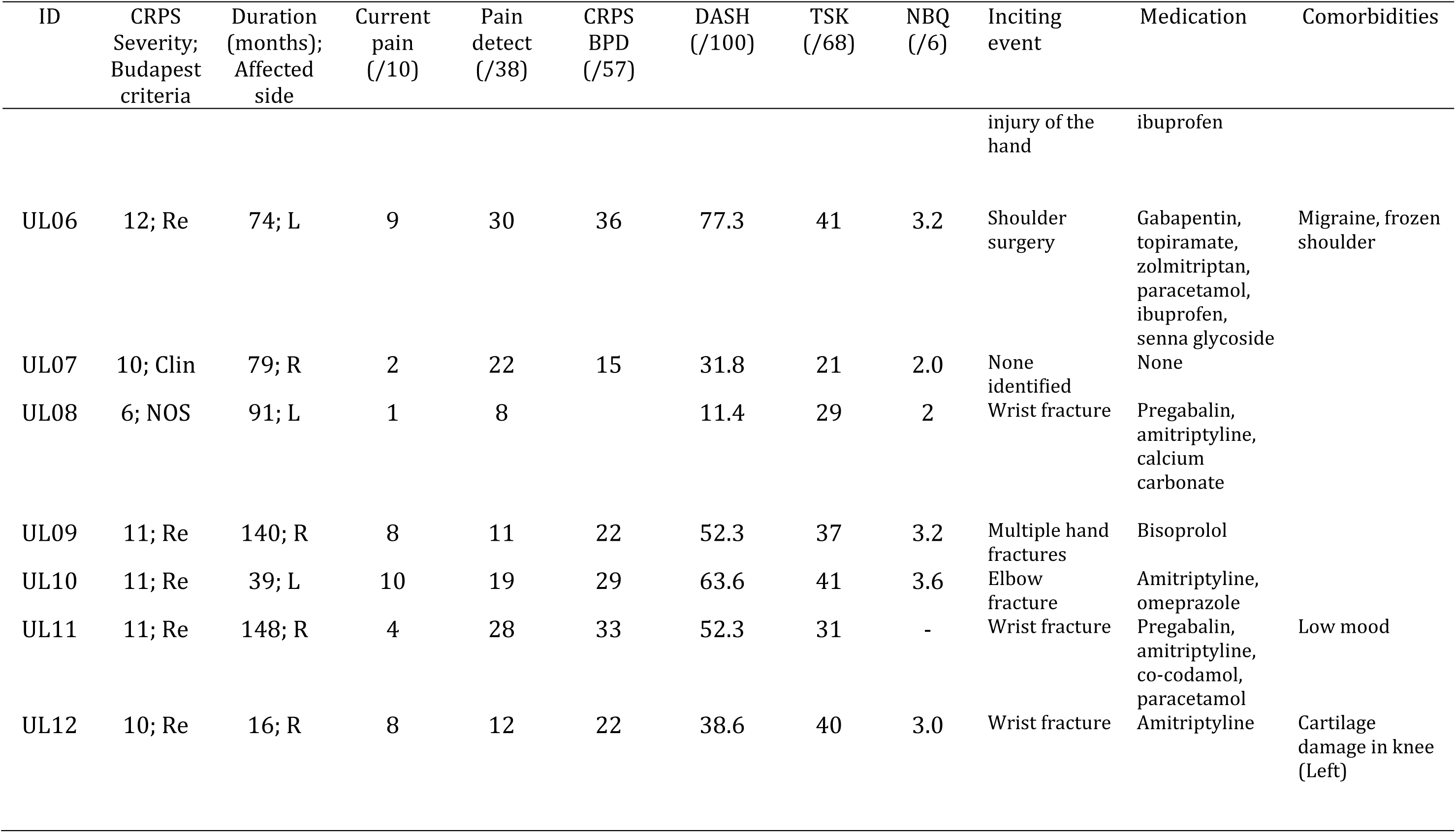
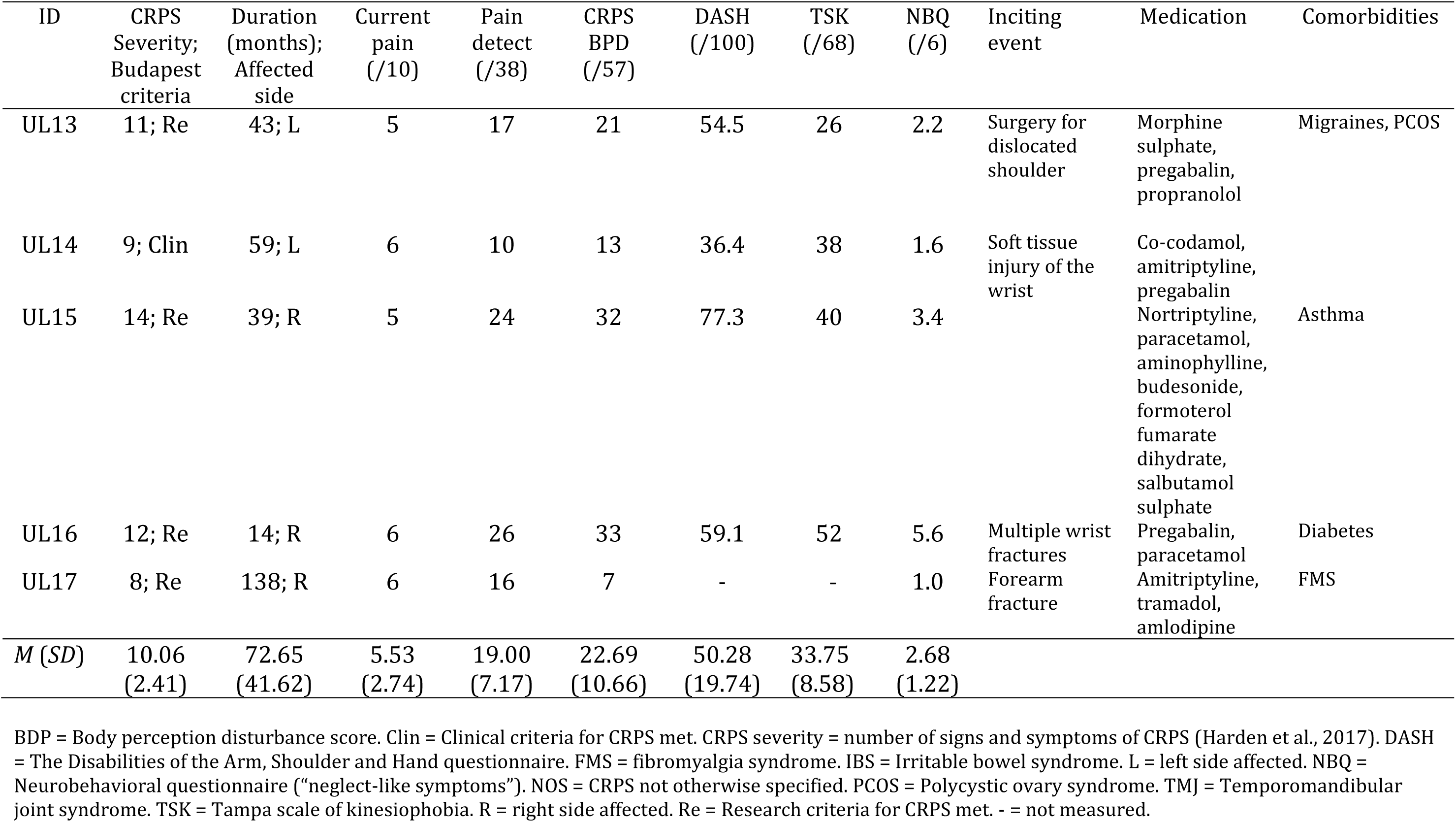
Clinical information for people with upper limb CRPS.

Participants were excluded if they reported a history of brain injury, brain disorders, or psychiatric disorders that can be associated with pronounced perceptual changes (e.g. schizophrenia; Tseng et al., 2015). Because the study involved exposure to a magnetic motion capture system, we also excluded people with a pacemaker, spinal cord stimulator or similar devices, and women who were pregnant or breastfeeding. All participants reported having normal or corrected-to-normal vision, and sufficient motor abilities to perform the movements required for the task. The study complied with the 2013 declaration of Helsinki and had ethical permission from the UK Health Research Authority (REC reference 12/SC/0557).

### 2.2. Stimuli and procedure

#### 2.2.1. Questionnaire measures

After providing informed written consent, participants completed questionnaire measures. All participants completed the Edinburgh handedness inventory (Oldfield, 1971), in which a negative score (<-40) indicates left-handedness, and a positive score (>40) indicates right-handedness. Three people with CRPS were classed as left-handed, four as ambidextrous, and eight as right-handed. Two control participants were classed as left-handed, three as ambidextrous, and 11 as right-handed.

People with CRPS completed additional questionnaires as follows (Table 1). Neuropathic components of pain were assessed by the pain DETECT questionnaire (Freynhagen, Baron, Gockel, & Tölle, 2006), where a score above 19/38 suggests that a neuropathic component is likely (>90% probability). The QuickDASH (Gummesson, Atroshi, & Ekdahl, 2003) was used to evaluate the degree of upper limb disability, where more severe disability is indicated by a higher score (/100). The Tampa Scale of Kinesiophobia (Miller, Kori, & Todd, 1991) was used to measure pain-related fear of movement and re-injury, where scores range from 17 (no kinesiophobia) to 68 (highest possible kinesiophobia). Body representation distortion was assessed by the Bath CRPS Body Perception Disturbance Scale (Lewis & McCabe, 2010), scored from zero (no body perception disturbance) to 57 (highest possible body perception disturbance).

Finally, severity of “neglect-like symptoms” was assessed by the Neurobehavioral questionnaire (Frettlöh, Hüppe, & Maier, 2006; Galer & Jensen, 1999), scored from one (no “neglect-like symptoms”) to six (highest possible severity of “neglect-like symptoms”).

#### 2.2.2. Prism adaptation

Participants performed a dynamic prism adaptation paradigm (Prablanc et al., 2019) that involved both open- and closed-loop trials (Figure 2). Because there has been some suggestion that adaptation to shifts towards the affected side might exacerbate pain for people with CRPS (Sumitani et al., 2007), we used adaptation to optical shifts away from the affected/non-dominant side (leftwards for 9/17 people with CRPS; leftwards for 3/18 controls). For the prism exposure trials, participants wore goggles fitted with Fresnel lenses that shifted vision laterally by 35 dioptres (∼19°). Participants completed the prism adaptation protocol with each hand in a randomised and counterbalanced order. During open-loop trials participants’ vision of their hand was occluded, and they performed pointing movements to a central visual target (i.e. 0°). For closed-loop trials, participants had vision of their hand when their arm was fully extended, whereas the arm from the wrist up was concealed (i.e. terminal exposure). Visual targets for closed-loop trials were 10° to the left or right of centre. For each hand, participants completed closed-loop and open-loop trials across five phases (see Figure 2). First, in the baseline phase, participants completed one block of 20 closed-loop trials and one block of 15 open-loop trials with unperturbed vision. Second, in the prism exposure phase, they performed 100 closed-loop trials towards targets viewed through the prism goggles, split into six blocks (Prism exposure – closed loop 1 (P-CL1), P-CL2, P-CL3, P-CL4, P-CL5, P-CL6). The blocks of closed-loop trials were separated by blocks of two open-loop trials (Prism exposure – open loop 1 (P-OL1), P-CL2, P-OL3, P-OL4, P-OL5) towards targets view through unperturbed vision. Third, in the after-effect phase, participants performed another block of 15 open-loop trials towards targets viewed with unperturbed vision. Fourth, in the washout phase they performed 60 closed-loop trials towards targets viewed with unperturbed vision, split into six blocks (Washout – closed loop 1 (W-CL1), W-CL2, W-CL3, W-CL4, W-CL5, W-CL6). The blocks of closed-loops trials were once again separated by blocks of two open-loop trials (Washout – open loop 1 (W-OL1), W-CL2, W-OL3, W-OL4, W-OL5) towards targets view through unperturbed vision. Finally, in the retention phase participants performed one block of 15 open-loop trials with unperturbed vision. The entire procedure was completed once for each hand, in a counterbalanced order. For each set of 10 closed-loop trials, five left targets and five right targets were presented, in a randomised order.

**Figure 2.**
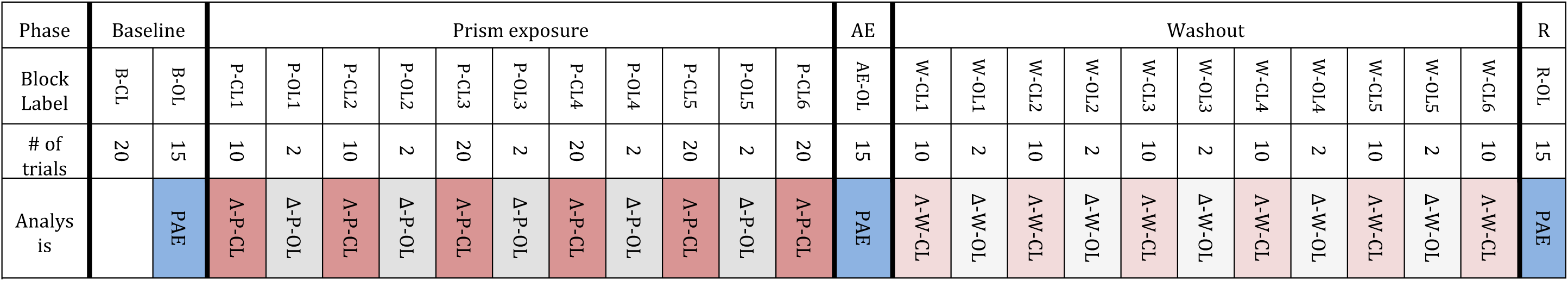
Prism adaptation protocol. Colours indicate what analysis each phase or block corresponds to. AE = after-effect; B = baseline; CL = closed loop pointing; OL = open-loop pointing; PAE = prism adaptation after-effects; PC = closed-loop trials during prism exposure phase; PO = open-loop trials during prism exposure phase; R = retention of after-effect; WC = closed-loop trials during washout phase; WO = open-loop trials during washout phase; A- = change in endpoint errors during open-loop trials; A = exponential decay of endpoint errors during closed-loop trials.

The first four people with CRPS to participate performed an additional 42 trials with each hand (two open-loop trials followed by 40 closed-loop trials) at the end of the washout phase. Despite their relatively good upper limb mobility, they found it difficult to complete all 287 pointing movements with their affected hand. We therefore reduced the number of trials in the washout phase for the remaining participants to that represented in Figure 2, and updated the preregistration accordingly (https://osf.io/6jpfg/). This resulted in a total of 245 trials per hand.

Participants were seated at a custom-built non-ferrous table (Fig. 3) with a Velcro-adjustable chin-rest affixed to the edge closest to the participant, and the magnet from the motion capture system attached to the underside of the table at the edge furthest away from the participant. A motion tracking sensor (7.9 mm × 7.9 mm × 19.8 mm) was placed on each of the participants’ index fingers with medical tape. To start a trial, participants placed their index finger on a raised tactile point (∼1 cm diameter) near to the trunk that was aligned with their body midline and the central target (the “start location”). The trial was programmed to start only if one of the sensors was first detected within ±2 cm laterally and ±3 cm distally of the start location. If a sensor was detected, a red light-emitting diode (LED) appeared in one of the target locations after a pause of 1 s. After a further 1 s, an audio cue (200 ms, 800 Hz) sounded. Participants were instructed that upon hearing the audio cue, they should point as quickly as possible with their fully extended arm and place their finger on the line leading to the visual target (Fig. 3). The visual target stayed illuminated for a further 3 s after the onset of the audio cue. Participants were instructed to bring their hand back to the start location upon the target extinguishing. The experimenter then pressed a computer key to allow the script to proceed to the next trial, which would once again start only if one of the sensors was detected within ±2 cm x ±3 cm of the start location. If interim open-loop trials (e.g. PO1, WO2) were completed incorrectly (e.g. a false start), they were repeated. No other trials were repeated.

**Figure 3.**
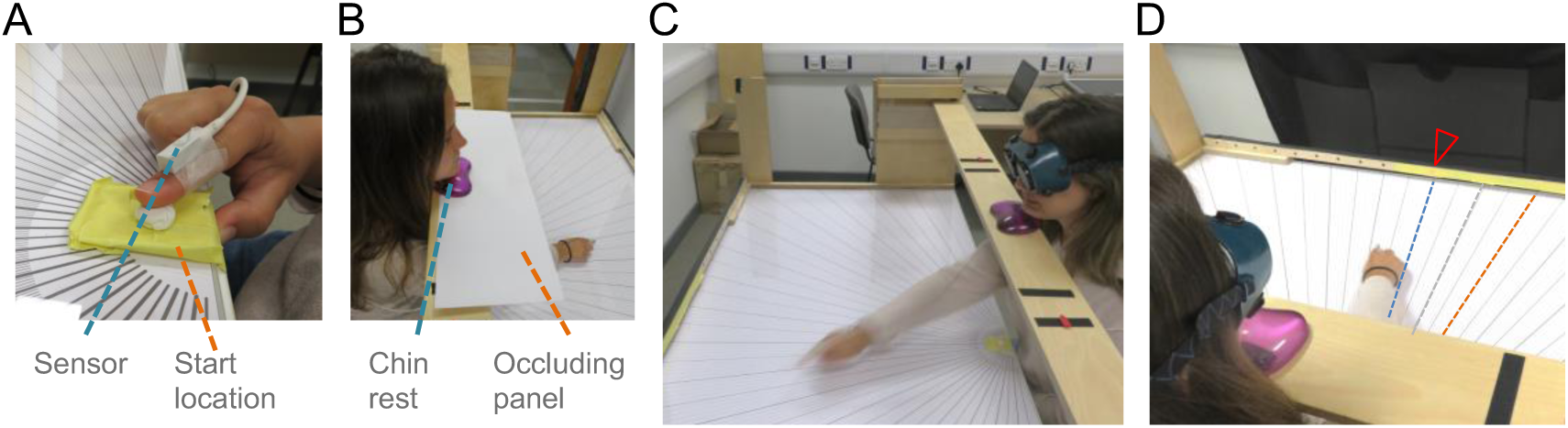
The prism adaptation table, chinrest, sensor (A), start location (A), and occluding panel (B). Panel B depicts an open-loop trial. Panels C and D depict closed-loop trials. The red arrow (D) indicates an example target location, with 10° left (blue), 10° right (orange), and central (i.e. 0°; grey) target axes superimposed.

We recorded 3D kinematic data at 240 Hz using an electromagnetic motion capture system (trakSTAR™, 3D Guidance^®^, Northern Digital Incorporated). The kinematic data was low-pass filtered using a second-order dual-pass Butterworth filter at 10 Hz. We calculated instantaneous velocities and accelerations by differentiating the data with a 5-point central finite difference algorithm, twice per axis. Velocity vectors were combined to yield resultant velocity, which was used to determine movement onset, and movement offset. The threshold for movement onset was set at 50 mm/s. Because people with CRPS can have motor impairments (e.g. spasms or arrests), we considered multiple criteria for movement offset for all participants: a threshold of 50 mm/s, and the point at which the sensor returned to the same vertical location as movement onset. The latter was taken as proxy of the hand being placed on the table. We visually inspected all trials (displacement, and resultant velocity plots), and manually adjusted movement onsets and movement offsets to satisfy the above criteria when needed. We deleted trials where a false start was detected (i.e. when movements faster than 50 mm/s were detected for the first sample; 2.52% of trials), and rotated movement trajectories to correct for a calibration error.

### 2.3. Statistical analyses and inference criteria

#### 2.3.1. Endpoint errors

Our primary analyses were of endpoint errors from closed-loop (3.2.) and open-loop (3.3.) trials. We calculated endpoint errors (°) as the angle between a two-dimensional straight line connecting the start location and movement offset, and a straight line from the start location to the target (i.e. the target axis). Endpoint errors made towards the affected/non-dominant side were expressed as negative values. We analysed trial level data using a linear mixed model regression, with participant ID and trial number as random effects.

#### 2.3.2. Exponential decay

For closed-loop trials during prism exposure phase (3.2.1.; P-CL1, P-CL2, P-CL3, P-CL4, P-CL5, P-CL6), and the washout phase (3.2.2.; W-CL1, W-CL2, W-CL3, W-CL4, W-CL5, W-CL6) we fitted exponential decay functions (*x* = *a* × e^*-b×n*^ + *c*) to endpoint errors for each person and each Hand (Facchin et al., 2018; Martin et al., 1996a; Nemanich & Earhart, 2015; O’Shea et al., 2014). We considered *x* as the endpoint error; *a* the initial error; *b* the decay constant; *c* the residual error; and *n* the trial number. The rate of error correction was expressed as the inverse of *b* (i.e. 1/*b*). The inverse of *b* equates to the half-life of the endpoint error reduction curve, and therefore indicates half the number of trials needed for endpoint errors to reach the asymptote (i.e. the residual error *c*).

To whether people with CRPS would show impaired strategic control relative to controls (Hypothesis 1), we compared the decay constant between Groups and the Hand used, using linear mixed models regression. We analysed the residual error (i.e. *c*) to address Hypothesis 2.

However, our primary analysis to address Hypothesis 2 (that people with CRPS would show impaired sensorimotor adaptation relative to controls) was to compare “raw” open-loop endpoint errors (2.3.1.) across the different experimental phases (Fig. 2). That is, we analysed endpoint errors without fitting an exponential decay function. We compared endpoint errors between Groups, Hand, and Phase (baseline, after-effect, retention). There were 15 open-loop pointing trials per phase for each hand.

To address Hypothesis 3, that sensorimotor realignment might develop and/or decline at a different rate for people with CRPS, compared to controls, we examined the development of the prism adaptation after-effects during prism exposure phase (3.3.2.), and their decline during the washout phase (3.3.3.). As with the primary analysis to address Hypothesis 2, we analysed endpoint errors without fitting an exponential decay function for these analyses. We performed separate analyses for the open-loop blocks during the prism exposure and washout phases (Fig. 2). For the prism adaptation phase, we compared endpoint errors between Groups, Hand, and PA Block (P-OL1, P-OL2, P-OL3, P-OL4, P-OL5); for the washout phase, we compared endpoint errors between Groups, Hand, and Washout Block (W-OL1, W-OL2, W-OL3, W-OL4, W-OL5).

#### 2.3.3. Trajectory orientations

We derived kinematic markers (3.4.) that have previously been associated with strategic recalibration (3.4.1.) and sensorimotor realignment (3.4.2.) during prism exposure (O’Shea et al., 2014). Specifically, we computed the tangential velocity vectors for peak acceleration (initial trajectory orientation) and peak deceleration (terminal trajectory orientation) and expressed them as the angle (°) relative to the target axis.

To address Hypothesis 4, that people with CRPS would show less evidence of trial-by-trial changes in movement plans to compensate for the prismatic shift than controls, we analysed the trial-by-trial change in initial trajectory orientations during early trials (i.e. closed loop trials 1 to 10 during prism exposure; P-CL1) for each Group, and Hand. These trial-by-trial changes allow for a specific strategy involved in strategic calibration to be examined (O’Shea et al., 2014), whilst eliminating any contribution from online corrections. We correlated the magnitude of endpoint errors on a given trial (*n*) during early prism exposure with the change in initial trajectory orientation on the subsequent trial (*n*+1). We performed this correlation on detrended data for only the early prism exposure trials, because error correction is typically only evident in these. To detrend the data, endpoint errors and initial trajectory orientations were fitted to the same exponential decay function as described in sections 2.3.2. (i.e. *x* = *a* × e^*-b×n*^ + *c*), and then the residuals were computed by subtracting the predicted values (i.e. *x*) from the observed value for each trial. The t-value for the correlation between these variables was calculated for each participant and each hand. If endpoint errors and initial trajectory orientations are unrelated then the *t*-values from these individual correlations should have a Gaussian distribution centred around zero. We aimed to test whether, for each Group and Hand, there was a linear relationship between endpoint errors on a given trial (*n*) and the change in movement plan on the next trial (*n*+1) by using one-sample *t*-tests to compare the individual participant *t*-values to zero. This analysis has been used previously to examine the changes in feedforward motor control during early prism exposure (O’Shea et al., 2014).

To further address Hypothesis 4, we performed unilateral Kolmogorov-Smirnov distribution tests (Vindras, Desmurget, & Baraduc, 2012) on the *p*-values obtained from individual correlations to see if they were biased towards zero. If individual *p*-values are biased towards zero it would provide further support for the presence of a linear relationship between endpoint errors on a given trial and the updated movement plan of the subsequent trial. This analysis therefore sheds light on the process of strategic error reduction during early prism exposure (O’Shea et al., 2014).

In our pre-registration we aimed to compare the terminal trajectory orientations between Groups, however this was not possible. Many participants made corrective finger movements in the later stage of a movement, once they became aware that they were about to miss the target. These late finger movements limited the information that could be derived from the point of peak deceleration (e.g. terminal trajectory orientations). When we filtered the data to remove these corrective finger movements the sample sizes were too small to make meaningful comparisons between Groups (CRPS n_affected_ = 8; CRPS n_non-affected_ =14; control n_non-dominant_ = 10; controls n_dominant_ = 15). Therefore, we do not report on the analyses of terminal trajectory orientations.

#### 2.3.4. Inference criteria

We processed and analysed the data in MATLAB (2018b; MathWorks, US), R (3.6.3; R Core Team, 2013), JAMOVI (1.1.9.0; The Jamovi Project, 2020), and JASP (0.12; JASP Team, 2018). Two people with CRPS were unable to complete the procedures with their affected hand. One of these participants (UL01) completed the full protocol with their non-affected hand. The other participant (UL15), however, was not able to do so due to the pain in her affected hand, and stopped after completing the third Open-loop Block in the washout phase (W-OL3) with her non-affected hand. As described in section 2.2.2., four participants with CRPS (UL01, UL03, UL11, UL13) completed additional trials in the washout phase. Because the number of washout trials can influence the measured retention of prism adaptation after-effects (Fernández-Ruiz & Díaz, 1999), we excluded the retention phase data of these participants from analyses related to after-effect retention.

We used linear mixed models regression for our main analyses, because this method allowed us to include all available data from all participants regardless of the missing elements just described. We also used mixed ANOVAs. To be concise, we only report interactions from analyses that address our hypotheses (i.e. that involve Group). We computed *p*-values from the linear mixed models regression with the Satterthwaite degrees of freedom (*lme4*, and *lmerTest* R packages), which has a lower Type-1 error rate than other methods (Luke, 2017). We report the ANOVA outputs from the linear mixed models analyses for ease of interpretation. We followed up any significant interactions with pairwise comparisons of the estimated marginal means derived from the linear mixed model (*emmeans*, and *multcomp* R packages), also using the Satterthwaite degrees of freedom. For these, and pairwise comparisons stemming from the ANOVAs, we applied Holm-Bonferroni corrections (Holm, 1979) for multiple comparisons, indicated by “*p*_adjusted_”. We considered *p*-values < .05 as statistically significant. Calculating effect sizes for linear mixed models is complicated by the challenges associated with estimating the degrees of freedom, and partitioning variance in linear mixed models (Rights & Sterba, 2019). 95% Confidence intervals are presented alongside effect size estimates, (Pek & Flora, 2018). For the follow-up tests of linear mixed models where we performed pairwise comparisons, we calculated Cohen’s *d* from the estimated marginal means, the population standard deviation estimated by the linear mixed model, and the Satterthwaite degrees of freedom (*emmeans* R package). For results that are reported in short, and as part of a cluster (e.g. *t*s(51) ≤ 2.42, *p*s_adjusted_ ≥ .340), the effect size estimate is reported using absolute values.

See preregistration for a full list of planned analyses (https://osf.io/6jpfg/). Analyses that were not preregistered are listed in *the Exploratory analyses* section (3.5.), or specified as exploratory.

## 3. Results

### 3.1. Summary statistics

Descriptive statistics for clinical data and questionnaire measures for people with CRPS are presented in Table 1. As prism adaptation can be influenced by the speed of movement (Redding et al., 2005), and its main outcome measures relate to the precision of pointing movements, we conducted two-way mixed ANOVAs on peak velocity, peak acceleration, and baseline closed-loop endpoint errors, with Group and Hand as factors. These revealed no meaningful differences between Groups or according to which Hand was used (Supplementary Text T1).

### 3.2. Closed-loop endpoint errors

#### 3.2.1. Exponential decay of endpoint errors during prism exposure

To test whether people with CRPS would show impaired strategic control relative to controls (Hypothesis 1), we analysed the strategic error reduction during closed-loop trials during the prism exposure phase (i.e. P-CL1, P-CL2, P-CL3, P-CL4, P-CL5, P-CL6; see Fig. 2).

Before analysing the constants derived from the fitted models, we analysed the model fit parameters. The model failed to converge, or there was no exponential fit for one person with CRPS (non-affected hand), and for one control (dominant hand). When we compared the model fit parameters the results suggested that the models were not found to differ across Groups and Hand (see Supplementary Text T2). Therefore, we proceeded to analyse the constants derived from the models (i.e. 1/*b*, and *c*; Fig. 4).

**Figure 4.**
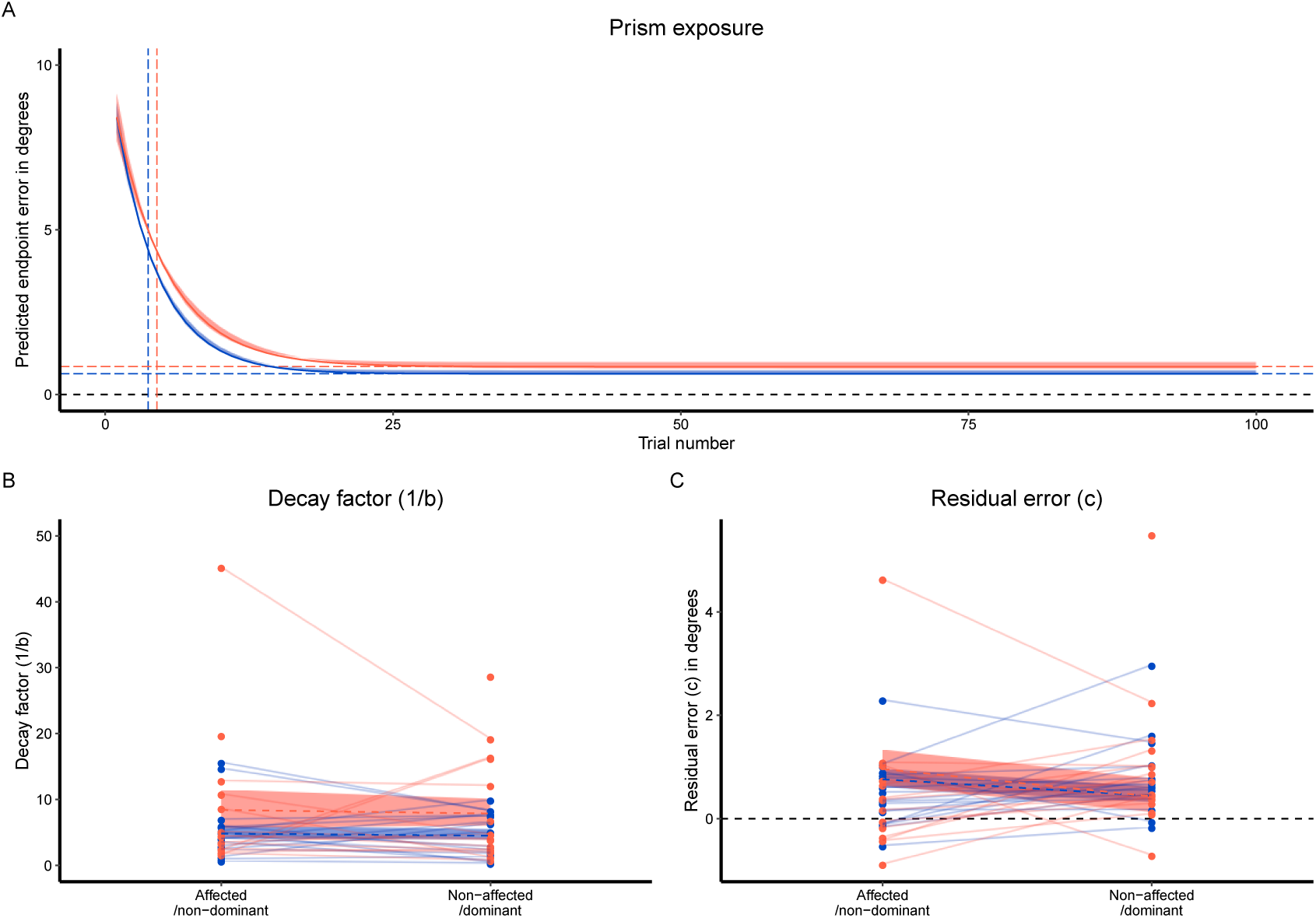
Exponential decay functions (*x* =*a*×*exp*^*-b×n*^+*c*) fitted to averaged data (A) and individual decay factors (B) and residual errors (C) for end-point errors for closed-loop trials during the prism exposure phase. In panel A, solid lines indicate the predicted value, and the boundaries of the shaded areas depict the 95% confidence interval for the constants (i.e. *a, b, c*) fitted to mean endpoint errors for each trial, split by Group. The black dashed lines (A, C) show perfect performance (i.e. zero degree error). Negative values indicate endpoint errors made towards the affected/non-dominant side (A, C). The coloured dashed lines (A) indicate the decay constant (i.e. 1/*b*; vertical lines), and the residual error (i.e. c; horizontal lines), for people with CPRS (red), and controls (blue). Points depict individual level data is presented for the decay factor (i.e. 1/b; B) and the residual error (i.e. c; C) for people with CRPS (red), and controls (blue). Data points are connected for each participant, given that they had data available and that we were able to fit it to an exponential decay function. The coloured dashed lines (B, C) indicate group means, and the boundaries of the coloured shaded areas show ± one Standard Error of the Mean.

The rate at which endpoint errors decayed did not differ between people with CRPS and controls, or between the affected/non-dominant and non-affected/dominant hand. That is, there was no significant difference between people with CRPS (*M*_*1/b*_ = 8.64, *95% CI* [5.39, 11.89]) and controls (*M*_*1/b*_ = 4.62, *95% CI* [1.50, 7.74]) on the decay factor (i.e. 1/*b*), *F*(1, 30.29) = 3.33, *p* = .078. There was also no significant main effect of Hand (affected/non-dominant *M*_1/b_ = 6.94, *95% CI* [4.38, 9.50]; non-affected/dominant *M*_*1/b*_ = 6.32, *95% CI* [3.77, 8.87]), *F*(1, 28.19) = 0.33, *p* = .629. Furthermore, there was no significant interaction between Group and Hand, *F*(1, 28.19) = 0.03, *p* = .860. These results therefore suggest that there was no difference in rate of endpoint error decay during prism exposure for people with CPRS and controls, while using either hand.

The residual endpoint error during prism exposure was not found to differ between people with CRPS and controls. That is, there was no significant difference in the residual error between people with CRPS (*M*_*c*_ = 0.75, *95% CI* [0.26, 1.25]) and controls (*M*_c_ = 0.61, *95% CI* [0.14, 1.08]), *F*(1, 26.29) = 0.19, *p* = .669. There was a tendency towards greater residual errors in the direction of the prismatic shift (i.e. towards the non-affected/dominant side) for the non-affected/dominant arm (*M*_*c*_ = 0.86, *95% CI* [0.48, 1.24]) compared to the affected/non-dominant arm (*M*_c_ = 0.50, *95% CI* [0.12, 0.88]), although not significant, *F*(1, 24.06) = 4.04, *p* = .056. There was no significant interaction between Group and Hand on the residual error, *F*(1, 24.05) < 0.01, *p* = .949. Our results therefore suggest that the residual error that remained once pointing correction had reached asymptote during prism exposure trials was not found to differ between people with CRPS and controls, and that there were no differences between groups that depended on the hand that was used.

#### 3.2.2. Exponential decay of endpoint errors during washout

To further test whether people with CRPS would show impaired strategic control relative to pain-free controls (Hypothesis 1), we analysed the strategic error reduction during closed-loop trials during the washout phase (i.e. W-CL1, W-CL2, W-CL3, W-CL4, W-CL5, W-CL6).

Prior to analysing the constants from the exponential decay function, we analysed the model fit. We were unable to fit an exponential decay function for two participants with CRPS, both for their affected hand. There were no clear difference in the model fits (i.e. RMSE, *adj. R*^*2*^; see Supplementary Text T2) between people with CRPS and controls, therefore we proceeded to analyse the constants derived from the models (i.e. 1/*b*, and *c*; Fig. 5).

**Figure 5.**
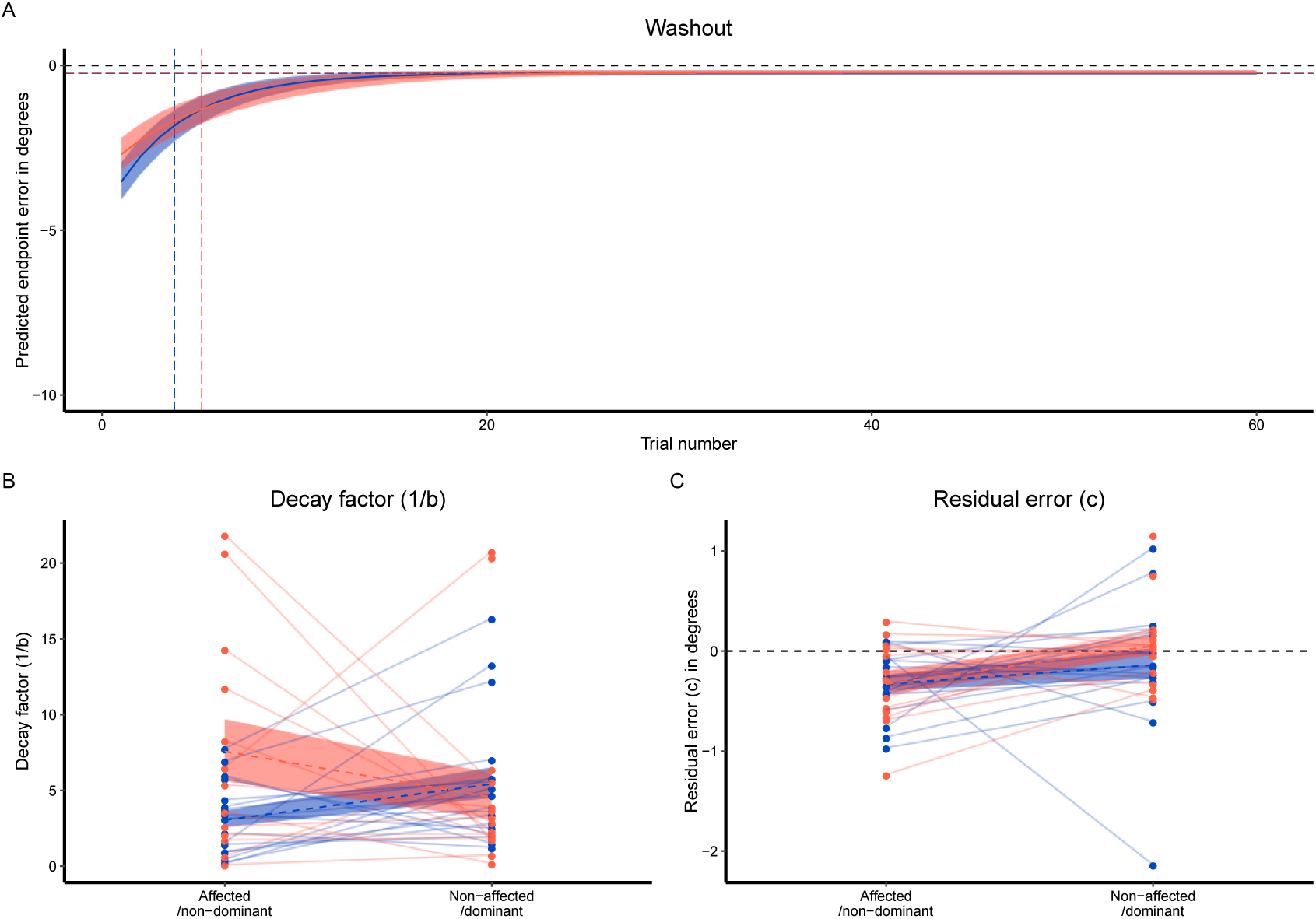
Exponential decay functions (*x* =*a*×*exp*^*-b×n*^+*c*) fitted to averaged data (A) and individual decay factors (B) and residual errors (C) for end-point errors for closed-loop trials during the washout phase In panel A, solid lines indicate the predicted value, and the boundaries of the shaded areas depict the 95% confidence interval for the constants (i.e. *a, b, c*) fitted to mean endpoint errors for each trial, split by Group. The black dashed lines (A, C) show perfect performance (i.e. zero degree error). Negative values indicate endpoint errors made towards the affected/non-dominant side (A, C). The coloured dashed lines (A) indicate the mean decay constant (i.e. 1/*b*; vertical lines), and the mean residual error (i.e. c; horizontal lines), for people with CPRS (red), and controls (blue). Points depict individual level data presented for the decay factor (i.e. 1/b; B) and the residual error (i.e. c; C) for people with CRPS (red), and controls (blue). Data points are connected for each participant, given that they had data available and that we were able to fit it to an exponential decay function. The coloured dashed lines (B, C) indicate group means, and the boundaries of the coloured shaded areas show ± one Standard Error of the Mean.

The rate at which closed-loop endpoint errors reduced during the washout phase was not found to differ between people with CRPS and controls, or for the affected/non-dominant and non-affected/dominant hand. That is, the decay rate did not significantly differ between people with CRPS (*M*_*1/b*_ = 6.09, *95% CI* [4.03, 8.15]) and controls (*M*_*1/b*_ = 4.23, *95% CI* [2.34, 6.12]), *F*(1, 28.69) = 1.85, *p* = .184. There was also no significant difference in decay rate that depended on the Hand (affected/non-dominant *M*_*1/b*_ = 5.34, *95% CI* [3.44, 7.23]; non-affected/dominant *M*_*1/b*_ = 4.98, *95% CI* [3.22, 6.74]), *F*(1, 28.12) = 0.09, *p* = .773.

There was a significant interaction between Group and Hand on the decay rate, *F*(1, 28.12) = 5.04, *p* = .033 (Fig. 5B). Although none of the follow-up comparisons were significant, the interaction appeared to be driven by a tendency for people with CRPS to have a larger decay factor when using the affected hand (*M*_*1/b*_ = 7.63, *95% CI* [4.75, 10.52]) than controls when using the non-dominant hand (*M*_*1/b*_ = 3.04, *95% CI* [0.58, 5.52]), *t*(51) = 2.42, *p*_adjusted_ = .340, *d* = 0.94, *95% CI* [0.14, 1.74]. The decay rates were numerically more similar, and the difference was in the opposite direction, for the non-affected/dominant hand (CRPS *M*_*1/b*_ = 4.54, *95% CI* [2.02, 7.07]; controls *M*_*1/b*_ = 5.42, *95% CI* [2.96, 7.87]), *t*(51) = 0.50, *p*_adjusted_ = .921, *d* = −0.18, *95% CI* [−0.90, 0.55]. This result indicates that there was a non-significant tendency for people with CRPS to need more trials to bring their endpoint errors back to baseline during washout when using their affected hand than controls when using their non-dominant hand.

The residual endpoint error during washout trials did not differ between people with CRPS and controls. That is, there was no main effect of Group (CRPS *M*_*c*_ = −0.14, *95% CI* [−0.32, 0.33]; controls *M*_*c*_ = −0.24, *95% CI* [−0.39, −0.08]) on residual errors (i.e. *c*), *F*(1, 62) = 0.62, *p* = .433. In contrast, the residual error was different between hands. Participants had a greater magnitude of residual error towards the affected/non-dominant side (i.e. the direction opposite to the prismatic shift) for their affected/non-dominant hand (*M*_*c*_ = −0.33, *95% CI* [−0.51, −0.16]) compared to their non-affected/dominant hand (*M*_*c*_ = −0.04, *95% CI* [−0.20, 0.12]), *F*(1, 62) = 6.09, *p* = .016. The interaction between Group and Hand on residual error was not significant, *F*(1, 62) = 0.63, *p* = .433. Therefore, our results provide no evidence of a difference in residual error during washout between people with CRPS and controls. Participants had greater residual error in the direction opposite to the prismatic shift for their affected/non-dominant hand, compared to their non-affected/dominant hand, which did not vary between Groups.

### 3.3. Open-loop endpoint errors

#### 3.3.1. Prism adaptation after-effects

To address the hypothesis that people with CRPS would show impaired sensorimotor adaptation relative to controls (Hypothesis 2), we analysed the effect of Group, Hand, and Phase (baseline, prism adaptation after-effects, retention) on endpoint errors during blocks of 15 open-loop trials (Fig. 6).

**Figure 6.**
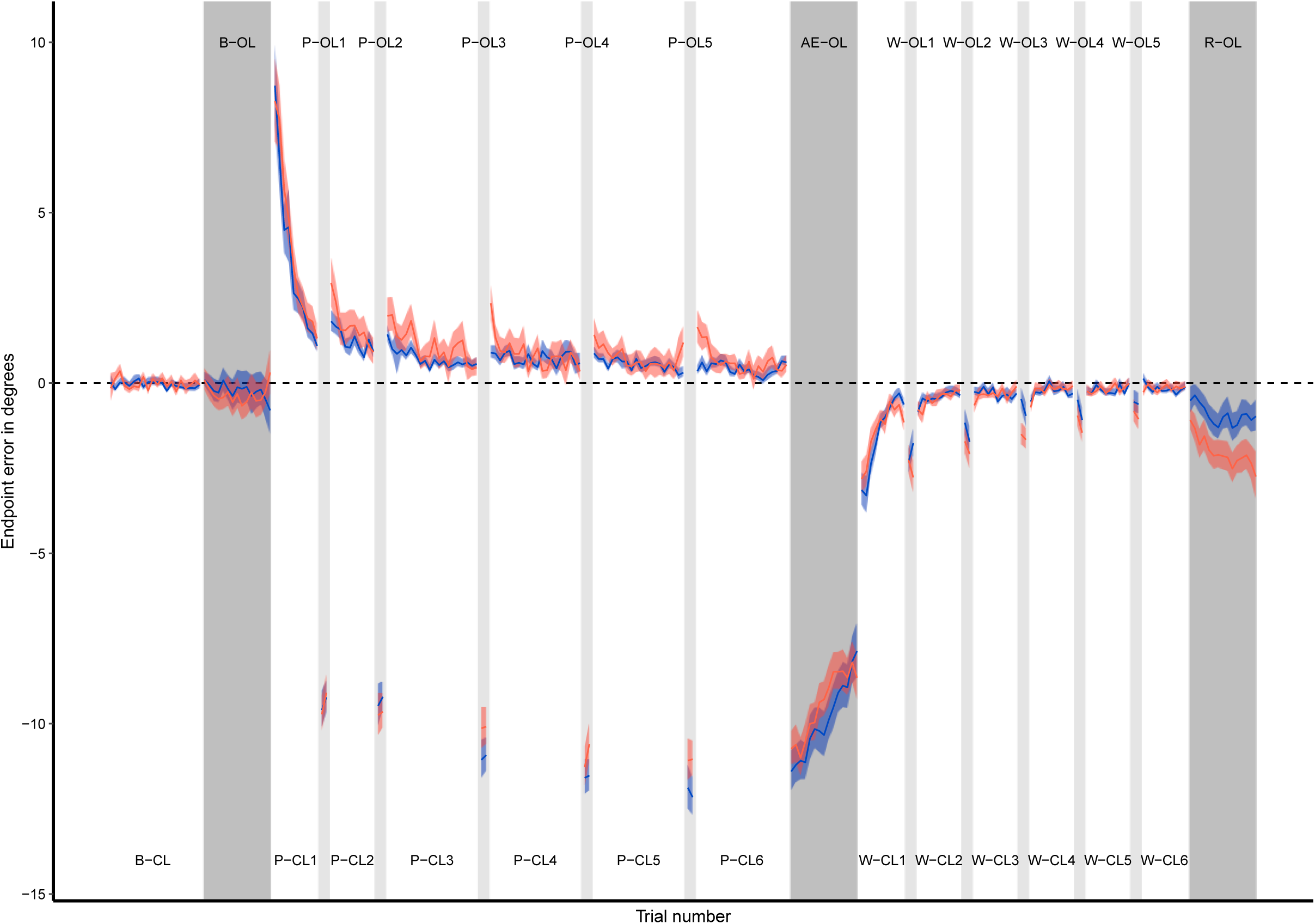
Endpoint errors in degrees are presented for people with CRPS (n = 17; red) and controls (n = 18; blue). The boundaries of the coloured shaded areas show the standard error of the mean. The grey shaded areas indicate open loop (OL) trials (i.e. when participants performed pointing movements without vision of their hand), for Open-loop Blocks (dark grey), Interim Open-loop Blocks (lighter grey), and Washout Open-loop Blocks (light grey). The white areas indicate closed-loop pointing, where participants had terminal exposure (i.e. they only had vision of their hand toward the end of a movement). The black dashed line shows the target orientation (i.e. zero degree error). Negative values indicate endpoint errors made towards the affected/non-dominant side. AE = after-effect phase; B = baseline phase; CL = closed-loop pointing; OL = open-loop pointing; P = prism adaptation phase; R = retention phase; W = washout phase

Participants adapted to the prismatic shift introduced by the goggles and showed some retention of this effect after the washout phase. There was a significant main effect of Phase, *F*(2, 42.93) = 1077.62, *p* < .001. Errors in the open loop trials performed directly after prism exposure were significantly deviated in the direction opposite to the prismatic shift (*M* = −9.77°, *95% CI* [−10.36, −9.17]) compared to both the baseline phase (*M* = −0.34°, *95% CI* [−0.93, 0.26]), *t*(43) = 42.61, *p*_adjusted_ ≤ .001, *d* = 4.07, *95% CI* [3.16, 4.96], and the retention phase (*M* = −1.46°, *95% CI* [−2.06, −0.86]), *t*(43) = 37.04, *p*_adjusted_ ≤ .001, *d* = −3.58, *95% CI* [−4.38, −2.78]. Endpoint errors were also significantly deviated for the retention phase compared to the baseline phase, *t*(43) = 5.01, *p*_adjusted_ < .001, *d* = 0.48, *95% CI* [0.26, 0.71]. These results indicate that adaptation to prismatic visual shifts produced sensorimotor after-effects (i.e. open-loop endpoint errors biased in the direction opposite to the prismatic shift relative to baseline), and that there was some retention of this effect after participants completed washout trials.

There was also a significant main effect of Hand on the open-loop endpoint errors, *F*(1, 2768.12) = 114.97, *p* < .001, whereby participants made greater errors towards their affected/non-dominant side (i.e. in the direction opposite to the prismatic shift) when using their affected/non-dominant hand (*M* = −4.33°, *95% CI* [−4.88, −3.78]) than with their non-affected/dominant hand (*M* = −3.38°, *95% CI* [−3.93, −2.83]). However, there was no main effect of Group (CRPS *M* = −4.00°, *95% CI* [−4.77, −3.23]; controls *M* = −3.71°, *95% CI* [−4.45, −2.96]), *F*(1, 32.63) = 0.32, *p* = .576.

There was a significant interaction between Group and Phase, *F*(2, 2766.60) = 21.21, *p* < .001. This interaction was superseded by a significant interaction between Group, Hand, and Phase (Fig. 7), *F*(2, 2762.70) = 22.37, *p* < .001. We followed up this interaction by performing four pairwise comparisons between each level of Group and Hand per Phase (baseline, prism adaptation after-effects, retention). For the baseline Phase, control participants made greater errors towards their non-dominant side with their non-dominant hand (*M* = −0.73°, *95% CI* [−1.54, 0.08]) compared to their dominant hand (*M* = 0.30°, *95% CI* [−0.52, 1.11]), *t*(950) = 5.04, *p*_adjusted_ < .001, *d* = 0.44, *95% CI* [0.27, 0.62]. There was no such difference for people with CRPS (affected *M* = −0.28°, *95% CI* [−1.12, 0.57]; non-affected *M* = −0.64°, *95% CI* [−1.47, 0.20]), and no significant differences between Groups for either Hand, *ts*(950) ≤ 1.70, *ps*_adjusted_ ≥ .663, *ds* ≤ 0.40. During the after-effect Phase, people with CRPS deviated further towards their affected side (i.e. the direction opposite to the prismatic shift) with their affected hand (*M* = −10.43°, *95% CI* [−11.27, −9.59]) than their non-affected hand (*M* = −8.70°, *95% CI* [−9.53, −7.86]), *t*(950) = 7.97, *p*_adjusted_ < .001, *d* = 0.75, *95% CI* [0.55, 0.94]. There was no significant difference between Hands for controls participants (non-dominant *M* = −10.17°, *95% CI* [−10.98, −9.35]; dominant *M* = −9.77°, *95% CI* [−10.59, −8.96]), and no differences between Groups for either hand, *ts*(950) ≤ 1.96, *ps*_adjusted_ ≥ .381, *ds* ≤ 0.46. In the retention phase, control participants made greater errors with their non-dominant hand (*M* = −1.83°, *95% CI* [−2.65, −1.02]) than their dominant hand (*M* = −0.05°, *95% CI* [−0.86, 0.78]), *t*(950) = 8.35, *p*_adjusted_ < .001, *d* = 0.77, *95% CI* [0.59, 0.95]). Similarly, people with CRPS made greater errors with their affected hand (*M* = −2.54°, *95% CI* [−3.40, −1.68]) than their non-affected hand (*M* = −1.42°, *95% CI* [−2.28, −0.55]), *t*(950) = 4.55, *p*_adjusted_ < .001, *d* = 0.49, *95% CI* [0.27, 0.70]. Comparing Groups, there was a tendency for people with CRPS to make greater errors in the direction opposite to the prismatic shift when using their non-affected hand than controls using their dominant hand. This difference was not significant after controlling for multiple comparisons (*t*(950) = 2.45, *p*_adjusted_ = .051), however the effect size was large (*d* = 0.59, *95% CI* [0.11, 1.07]). There was no significant difference between Groups when using their affected/non-dominant hand (*t*(950) = 1.28, *p*_adjusted_ = .263), with a medium effect size (*d* = 0.31, *95% CI* [−0.18, 0.79]). These results suggest that each Group had a tendency for larger (i.e., more negative) endpoint errors for their affected/non-dominant hand than their non-affected/dominant hand in at least two phases, and that there was a weaker evidence for this tendency for people with CRPS (compared to controls) to have larger endpoint errors persisting through the retention phase when using their non-affected/dominant hand.

**Figure 7.**
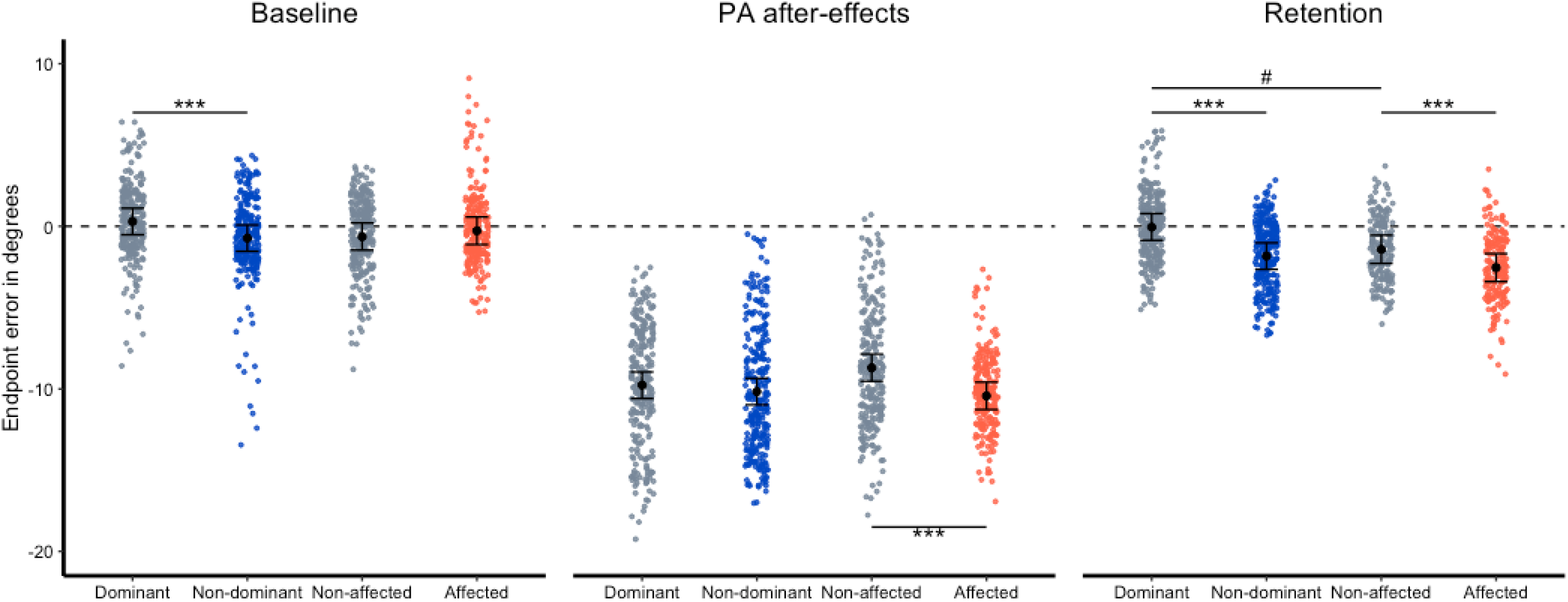
Open-loop endpoint errors for the baseline, after-effect, and retention phases are presented for people with CRPS (n = 17; red and grey) and controls (n = 18; blue and grey), for the affected (red)/non-dominant (blue) and the non-affected/dominant (grey) hands. Endpoint errors (in degrees) are presented for individual trials included in the linear mixed models regression analysis (i.e. for the 15 open-loop trials for each Open-loop Block, for each Hand, and for each participant). Black points and error bars depict the estimated marginal means and 95% confidence intervals derived from the analysis. The black dashed line shows the target orientation (i.e. zero degree error). Negative values indicate endpoint errors made towards the affected/non-dominant side. PA = prism adaptation. # *p*_adjusted_ = .051, *** *p*_adjusted_ < .001

To confirm that these results weren’t influenced by proprioceptive abilities, the direction of the prismatic shift, and/or the counterbalancing order, we ran three exploratory ANCOVAs with proprioceptive abilities as a covariate, comparing between the direction of prismatic shift and/or the counter balancing order. We found no evidence to suggest that our results were influenced by any differences between participants in proprioceptive abilities, *Fs*(2, 62) ≤ 1.03, *ps* ≥ .362, ƞ^2^ _p_ ≤ .03, the direction of the prismatic shift, *Fs*(2, 62) ≤ 0.62, *ps* ≥ .544, ƞ^2^ _p_≤ .02, or the counterbalancing order *Fs*(2, 62) ≤ 2.64, *ps* ≥ .080, ƞ^2^ _p_≤ .08 (Supplementary Text T3).

#### 3.3.2. Development of prism adaptation after-effects

To address the hypothesis that sensorimotor realignment might develop at a different rate for people with CRPS, compared to controls (Hypothesis 3), we analysed the effect of Group and Hand on endpoint errors during the Open-loop Blocks of the prism exposure phase (Fig. 6; P-OL1, P-OL2, P-OL3, P-OL4, P-OL5).

There was a main effect of Open-loop Block on endpoint errors, *F*(4, 617.25) = 23.50, *p* < .001. We followed this effect up by comparing each Interim Open-loop Blocks to the first block (i.e. PO1). This effect was driven by a greater magnitude of endpoint errors made during the fourth (PO4; *M* = −11.38°, *95% CI* [−12.30, −10.50]; *t*(5) = 6.52, *p*_adjusted_ = .018, *d* = 0.80, *95% CI* [0.24, 1.37]), and fifth (PO5; *M* = −11.65°, *95% CI* [−12.50, −10.77]; *t*(5) = 7.48, *p*_adjusted_ = .015, *d* = 0.92, *95% CI* [0.28, 1.55]) blocks compared to the first block (PO1; *M* = −9.50°, *95% CI* [−10.40, −8.62]). The second (PO2; *M* = −9.62°, *95% CI* [−10.50, −8.74]; *t*(5) = 0.70, *p*_adjusted_ = .704, *d* = 0.05, *95% CI* [−0.20, 0.30]), and third blocks (PO3; *M* = −10.60°, *95% CI* [−11.50, −9.72]; *t*(5) = 3.81, *p*_adjusted_ = .094, *d* = 0.47, *95% CI* [0.08, 0.86]) were not significantly different to the first block (i.e. PO1).

There was also a significant main effect of Hand, *F*(1, 620.79) = 8.32, *p* = .003, whereby participants made greater errors towards the affected/non-dominant side (i.e. in the direction opposite to the prismatic shift) when using their affected/non-dominant hand (*M* = −10.80°, *95% CI* [−11.70, −9.99]) compared to their non-affected/dominant hand (*M* = −10.30°, *95% CI* [−11.10, −9.45]. There was no significant main effect of Group, and no significant interactions *Fs*(4, 620.79) ≤ 2.27, *p* ≥ .060. These findings provide no evidence of a difference for development of prism-adaptation after-effects between people with CRPS and controls, and that this did not vary depending on the hand being used.

#### 3.3.3. Decline of sensorimotor after-effects

To address the hypothesis that sensorimotor realignment might decline at a different rate for people with CRPS, compared to controls (Hypothesis 3), we analysed the effect of Group and Hand on endpoint errors during the Open-loop Blocks of the washout phase (Fig. 6; W-OL1, W-OL2, W-OL3, W-OL4, W-OL5).

The prism adaptation after-effects decayed during washout trials. This was evidenced by a main effect of Washout Open-loop Blocks on endpoint errors, *F*(4, 5.04) = 10.92, *p* = .011. We followed this effect up by comparing each Washout Open-loop Block to the first block (i.e. WO1). This effect was driven by a greater magnitude of endpoint errors made during the fifth block (WO5; *M* = −0.76°, *95% CI* [−1.33, −0.18]) than the first block (WO1; *M* = −2.30°, *95% CI* [−2.88, −1.72]), *t*(5) = 4.97, *p*_adjusted_ = .044, *d* = −0.94, *95% CI* [−1.65, −0.23]. The second (WO2; *M* = −1.70°, *95% CI* [−2.27, −1.12]; *t*(5) = 2.28, *p*_adjusted_ = .217, *d* = −0.37, *95% CI* [−0.79, 0.05]), third (WO3; *M* = −1.20°, *95% CI* [−1.77, −0.62]; *t*(5) = 5.84, *p*_adjusted_ = .077, *d* = −0.68, *95% CI* [−1.24, −0.11]), and fourth (WO4; *M* = −0.98°, *95% CI* [−1.56, −0.41], *t*(5) = 4.73, *p*_adjusted_ = .061, *d* = −0.80, *95% CI* [−1.43, −0.17]) blocks were not significantly different to the first block (i.e. WO1) after correcting for multiple comparisons.

Participants had a greater retention of prism adaptation after-effects for their affected/non-dominant hand. That is, there was a significant main effect of Hand on endpoint errors during Washout Open-loop Blocks, *F*(1, 618.86) = 93.92, *p* < .001. Participants made greater errors in the direction opposite to the prismatic shift (i.e. towards the affected/non-dominant side) when using their affected/non-dominant hand (*M* = −2.01°, *95% CI* [−2.47, −1.54]) compared to their non-affected/dominant hand (*M* = −0.77°, *95% CI* [−1.23, −0.30]). There was no significant main effect of Group, and no significant interactions that involved Group, Hand, and/or Washout Open-loop Block, *F*s(4, 618.86) ≤ 1.63, *p*s ≥ .211. These findings suggest that the decay of prism adaptation after-effects did not differ between people with CRPS and controls, and did not vary depending on the Hand being used.

### 3.4. Kinematic changes during prism exposure

To address the hypothesis that people with CRPS would show less evidence of trial-by-trial changes in movement plans to compensate for the prismatic shift than controls (Hypothesis 4), we analysed the change in initial trajectory orientations during early trials (i.e. closed loop trials 1 to 10 during prism exposure; P-CL1) for each Group, and Hand.

Prior to analysing the detrended data for initial trajectory orientations, we inspected the model fit. We were unable to fit initial trajectory orientations to the exponential decay function for one person with CRPS (non-affected hand), and two controls (one non-dominant hand, one dominant hand). When we compared the model fit parameters for the remaining participants, the amount of variance explained by the models did not differ between people with CRPS and controls, or depending on the Hand used (Supplementary Text T2).

There was no evidence for trial-by-trial feedforward control for controls or people with CRPS, from either the analysis of t-values or p values. That is, we did not find any evidence of a significant linear relationship between endpoint errors and changes in initial trajectory orientation for either Group using either Hand, *t*s(14) ≤ 1.43, *p*s_adjusted_ ≥ .174, *d*s ≤ 0.36. When we pooled the data for each Group, and explored it for each hand, we found evidence that t-values were different from zero for the non-affected/dominant hand (Supplementary Text T4). The analysis of *p*-values from individual correlations showed no evidence of a bias towards zero for either Hand (affected, non-affected, dominant, non-dominant), *D*^+^ ≤ 0.18, *ps* ≥ .104. Therefore, only the data pooled across all participants provides evidence that trial-by-trial feedforward motor control was used to reduce endpoint errors for the non-affected/dominant hand during early prism exposure trials. However, there was no such evidence when data were considered separately for each Group from either *t*-values or from *p*-values.

### 3.5. Exploratory analyses

To further test idea that CRPS symptoms might be related to impaired strategic control and/or sensorimotor realignment, we analysed correlations for each Group (Figures S1 and S2). For control participants the variables that were included for each hand were peak velocity as a measure of movement kinematics, as well as the following variables calculated for the prism adaptation and washout phases: differences in open-loop endpoint errors from the baseline phase; and decay factor (1/b) and residual error (c) for closed-loop endpoint errors. For people with CRPS, we additionally included variables for clinical characteristics (CRPS severity, CRPS duration, baseline pain) and questionnaire measures (neuropathic type pain, fear of movement, upper limb disability, body perception disturbance, “neglect-like symptoms”). Below we report on the correlations that involve clinical variables and prism adaptation outcomes for people with CRPS. The other correlations are reported in the supplementary materials (Supplementary Text T3).

There were no significant correlations between any of the clinical characteristics and differences in open-loop endpoint errors for the exposure or washout phases compared to baseline for either hand. However, baseline pain showed a positive correlation with the decay factor for the affected hand during prism exposure (i.e. 1/*b*), *r* = .52, *p* = .046, which suggests that those experiencing more pain also needed more closed-loop trials for endpoint errors to decay. Furthermore, CRPS severity showed large positive correlations with the residual error (i.e. *c*) for the non-affected hand during prism exposure and washout trials, *r*s ≥ .54, *p*s ≤ .030. More positive residual errors indicate weaker sensorimotor realignment in the prism exposure phase (see Figure 3), and weaker retention of prism after-effect in the washout-phase (see Figure 4). Thus, these correlations suggest that more severe CRPS is associated with lesser sensorimotor realignment when using the non-affected hand, although the direction of this association may vary between parameters. Similarly, upper limb disability showed strong, positive correlations with the residual errors for each hand during prism exposure and washout, *r*s ≥ .57, *p*s ≤ .020. Our exploratory correlations therefore provide some evidence that CRPS severity and upper limb disability were related to sensorimotor processing, although only for some analysed variables.

## 4. Discussion

Our study was the first to characterise sensorimotor adaptation in people with CRPS. The results did not support our main hypotheses, as we found no evidence that prism adaptation is impaired for people with CRPS. Instead, we found some indications that CRPS might lead to a greater propensity for sensorimotor realignment, because people with CRPS showed a larger magnitude of open-loop pointing errors for their affected hand than their non-affected hand after prism exposure, whereas there was no difference between hands for the controls. Generally, however, we found no differences between people with CRPS and controls on most prism adaptation measures. Below we discuss the findings related to strategic control (4.1.), sensorimotor realignment (4.2.), and the retention of prism adaption after-effects (4.3.). We also consider the theoretical (4.4.1.) and clinical (4.4.2.) implications of our findings, and potential limitations (4.5.) of our study.

### 4.1. Strategic control

We found no evidence to suggest that strategic recalibration was disrupted for people with CRPS (Hypothesis 1). That is, we did not observe any difference between people with CRPS and controls on endpoint errors during prism exposure, or during washout. We also did not find any group differences in the number of trials needed for endpoint errors to decay during prism exposure.

### 4.2. Sensorimotor realignment

Overall, we found very little evidence that sensorimotor realignment was impaired for people with CRPS compared to controls (Hypothesis 2). We did not find any difference in the residual errors during prism exposure or during washout. There was no overall difference in the endpoint errors made during open-loop pointing directly after prism exposure. However, when the hands were considered separately, people with CRPS made greater endpoint errors in the prism adaptation after-effect phase with their affected hand than their non-affected hand, whereas there was no difference between hands for the controls in this phase. This finding is somewhat tentative, because direct comparisons between the groups revealed no differences in the open-loop pointing errors for the affected versus non-dominant and non-affected versus dominant hands. Nonetheless, these results suggest that, if anything, sensorimotor realignment was more pronounced for the affected hand, directly contradicting the idea that any impairment in CRPS would be more/only evident when using this hand (Hypothesis 5).

More pronounced sensorimotor realignment in the affected hand could relate to lower stability of representations of the body and space. In a previous study, we showed that following tool use, people with upper limb CRPS had more pronounced changes in their arm and peripersonal space representations compared to matched controls (Vittersø et al., 2020). Furthermore, in people with upper limb CRPS, representations of the affected and non-affected arms updated differently. Proprioceptive reference frames are closely linked to bodily representations (Medina & Coslett, 2010), and are updated during sensorimotor realignment (Jeannerod & Rossetti, 1993; Redding et al., 2005). Thus, the findings of the current study could be interpreted as further evidence that representations of the body and peripersonal space are less stable- or more malleable – in CRPS.

An outstanding issue if what mechanisms underly this malleability. One possibility is that it could relate to impaired online control due to the motor deficits that are diagnostic of CRPS. Differences in sensorimotor realignment between hands would be expected if participants used online control to different extents for each hand when reducing endpoint error during prism exposure (Redding et al., 2005). Specifically, greater post prism adaptation open-loop pointing errors we observed for the affected hand (compared to the unaffected hand) suggests that participants made less use of strategic correction with that hand, allowing for a greater proportion of error reduction to occur through sensorimotor realignment. However, we found no evidence of any difference in strategic control between hands for the CRPS group, or between the participants with CRPS and the controls (4.1). Therefore, our data do not support this explanation.

### 4.3. Retention of prism adaptation after-effects

The decay of the prism adaptation after-effects was not found to differ between groups during open-loop washout trials. There was also no difference in the residual error during closed-loop washout trials between groups. There was a tendency for people with CRPS to retain prism adaptation after-effects for longer and/or to a greater extent than controls, when using their non-affected/dominant hand, although this effect was not significant. Nonetheless, this pattern is worth noting because such a difference, if significant, would suggest that people with CPRS have more rigid sensorimotor representations of their non-affected hand. This is consistent with our finding that they showed a smaller magnitude of post prism adaptaiton open-loop errors for their non-affected hand than their affected hand. However, we did not find any evidence to suggest that the retention of prism adaptation after-effects was impaired for people with CRPS compared to controls.

### 4.4. Implications

#### 4.4.1. Theoretical implications

Sensorimotor incongruence has been theorised to contribute to the maintenance of certain pathological pain conditions (Harris, 1999), such as CRPS (McCabe & Blake, 2007). One of the assumptions implicit to this theory is that people with CRPS and related conditions do not resolve such conflict by adapting to incongruent sensory and motor information. Under normal conditions, sensorimotor adaptation would occur to compensate for discrepant sensory and motor information (Wolpert et al., 2011) if the source of the incongruent information is assumed to be the same (Wei & Kording, 2009). In contrast to theoretical predictions (Harris, 1999), we did not observe any impairment for people with CRPS in the magnitude of sensorimotor adaptation following exposure to a lateral visual distortion, nor in the rate at which the after-effect developed across exposure blocks. In fact, we observed some tentative indications that the painful limb has an enhanced propensity for sensorimotor realignment. If pain is a result of conflicting sensorimotor information, then these finding suggests that this conflict is not driven by impaired sensorimotor adaptation for people with CRPS. We also did not find any correlations between the magnitude of sensorimotor realignment and clinical characteristics for people with CRPS. Therefore, our results suggest that sensorimotor integration is not impaired for people with CRPS, and that it is not related to pain and physical symptoms. These findings contradict the implicit assumption of the sensorimotor theory of pain, or at least suggest that sensorimotor conflict is not due to any impairment in adaptation.

Despite the cortical origin of pain proposed by the sensorimotor theory of pain, we did not find any evidence indicative of disrupted cortical processing in CRPS, as strategic recalibration, sensorimotor realignment, and its retention were not impaired. Strategic recalibration is dependent on cerebello-parietal processing (Panico et al., 2020). Our findings therefore suggest that these networks are not significantly altered in people with CRPS. Similarly, our findings provide no evidence to suggest that sensorimotor realignment during prism exposure is altered in CRPS, as would be if cerebellar processing, motor cortical processing(Panico et al., 2020), and/or the connectivity between these regions (Tsujimoto et al., 2019) were disrupted. Furthermore, our findings provide no evidence of altered M1 activation (O’Shea et al., 2017; Panico et al., 2017) for people with CRPS, as people with CRPS did not consistently retain greater prism adaption after-effects than controls. Therefore, our findings do not support the idea that cortical processing is disrupted for people with CRPS.

#### 4.4.2. Clinical implications

Our findings have implications for the application of prism adaptation as a treatment for CRPS. Several studies have examined the efficacy of treating CRPS with prism adaptation (Bultitude & Rafal, 2010; Christophe et al., 2016; Halicka, Vittersø, et al., 2020b; Sumitani et al., 2007). The first report of its application in CRPS suggested that the benefits were due to a normalisation of attention biases, or that pain relief might be due to improving sensorimotor integration (Sumitani et al., 2007). However, attention biases are not always found for people with CRPS (e.g. De Paepe et al., 2020; Filbrich et al., 2017; Halicka, Vittersø, et al., 2020a), and the therapeutic benefits have been reported in the absence of any consistent bias (Christophe et al., 2016). Furthermore, our findings suggest that people with CRPS do not have difficulties with adapting to the prism goggles. The therapeutic benefits of treating CRPS with prism adaptation are therefore unlikely to be due to correcting attention biases, or improving sensorimotor integration. However, our findings also suggest that any lack of an improvement is not likely explained by difficulties adapting to prism goggles (e.g. due to impaired sensorimotor integration). Taken together, these findings may shed light why the largest trial to date found no benefit of prism adaptation compared to a sham treatment (Halicka, Vittersø, et al., 2020b). Our findings are therefore compatible with recent evidence in questioning the efficacy of prism adaptation for treating CRPS.

### 4.5. Limitations

Fourteen of the people with CRPS had previous experience with prism adaptation as part of a double-blind randomized controlled trial, an average of 15 months prior to participating in our study. Previous experience with prism adaptation could have reduced the magnitude of adaptation (Martin, Keating, Goodkin, Bastian, & Thach, 1996b). The two studies took place over a year apart, and the participants had not yet been unblinded as to which treatment (real or sham prism adaptation) they had received in the previous study. For these reasons, it is unlikely that there were any additive effects of previous prism exposure, although we cannot rule out that participants remembered a strategy for compensating for the lateral optical distortion. In this case, our current findings would underestimate the sensorimotor after-effects in CRPS. Therefore, our results would still not support the hypothesised impairment in sensorimotor realignment.

## 5. Conclusions

Our study was the first to characterise sensorimotor adaptation in people with CRPS. Using prism adaptation, we found no evidence of impaired strategic recalibration or sensorimotor realignment for people with CRPS. People with CRPS showed greater prism adaptation after-effects for their affected hand than for their non-affected hand. This contradicts our hypothesis that any deficit would be more pronounced when using the affected limb, and instead adds to previous evidence that representations of the body and peripersonal space are more pronounced in CRPS. Overall, our findings indicate that people with CRPS were able to correct for the optical displacement introduced by the prisms, and that their sensorimotor system adapted appropriately to this displacement. These findings challenge existing theories of how pain might be maintained in the absence of clear tissue pathology, and add to our understanding of neuropsychological changes in CRPS.

## Acknowledgements

The authors thank Ms Eve Evans for her help with the study, Prof Andrea Serino and Dr Karin Petrini for their insightful comments on the manuscript, and the participants for volunteering their time.

## 6. Supplementary material

### 6.1. Supplementary Figure S1: Exploratory correlations – CRPS

**Figure S1.**
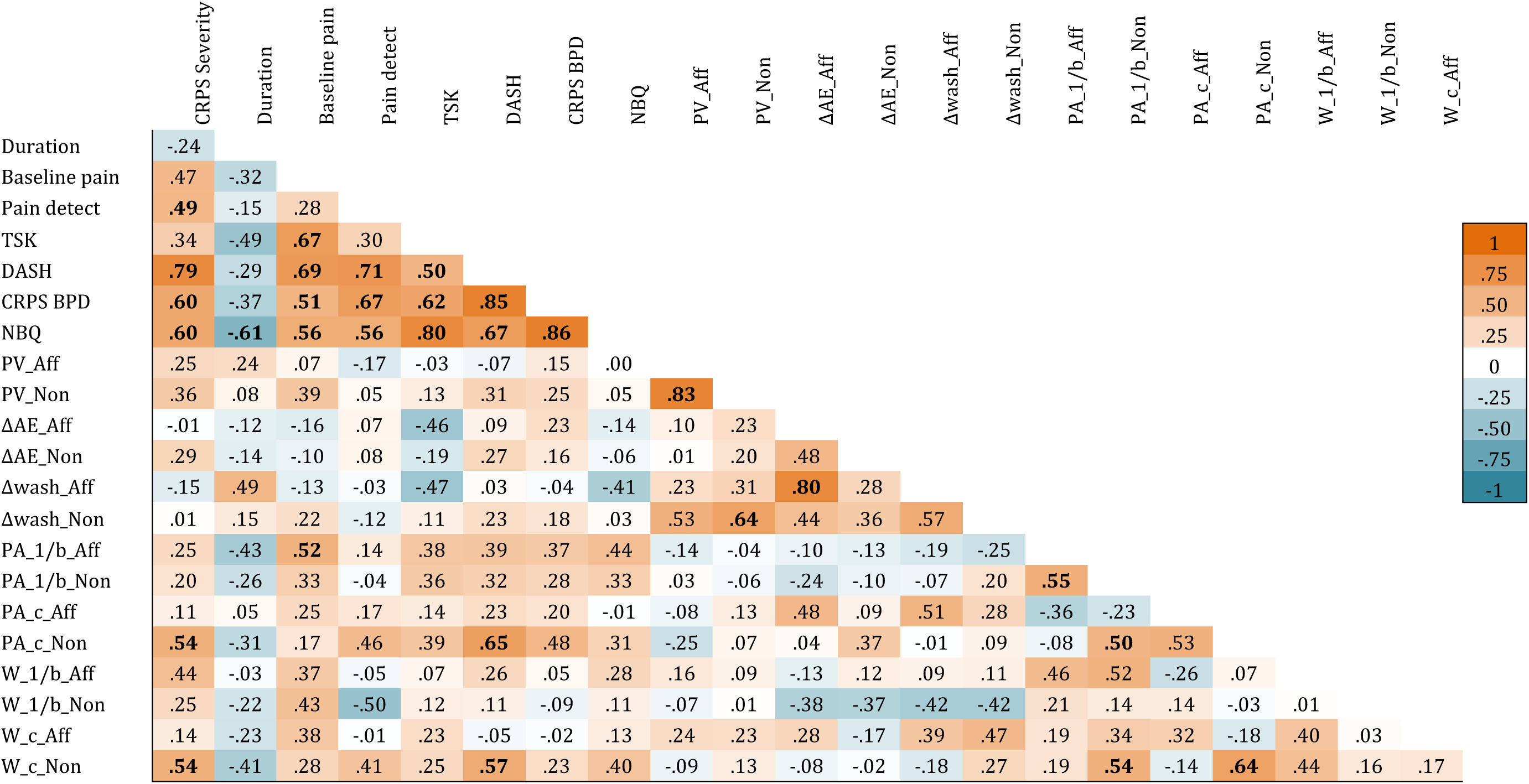
Pearson correlation matrix for people with upper limb CRPS (n = 17). Significant correlations (i.e. *p* < .05) are presented in boldface. Aff = CRPS affected limb; DASH = QuickDASH (Gummesson, Atroshi, & Ekdahl, 2003); ΔAE = endpoint errors from prism adaptation after-effects Open-loop Block, after subtracting baseline pointing error; Δwash = endpoint errors from retention Open-loop Block, after subtracting baseline pointing error; Duration = CRPS duration in months; CRPS BPD = Bath CRPS Body Perception Disturbance Scale (Lewis & McCabe, 2010); PV = peak velocity; TSK = Tampa Scale of Kinesiophobia; NBQ = Neurobehavioral questionnaire (Frettlöh et al., 2006; Galer & Jensen, 1999); Non = Non-affected limb; PA = Prism adaptation; W = Washout.

### 6.2. Supplementary Figure S2 Exploratory correlations - controls

**Figure S2.**
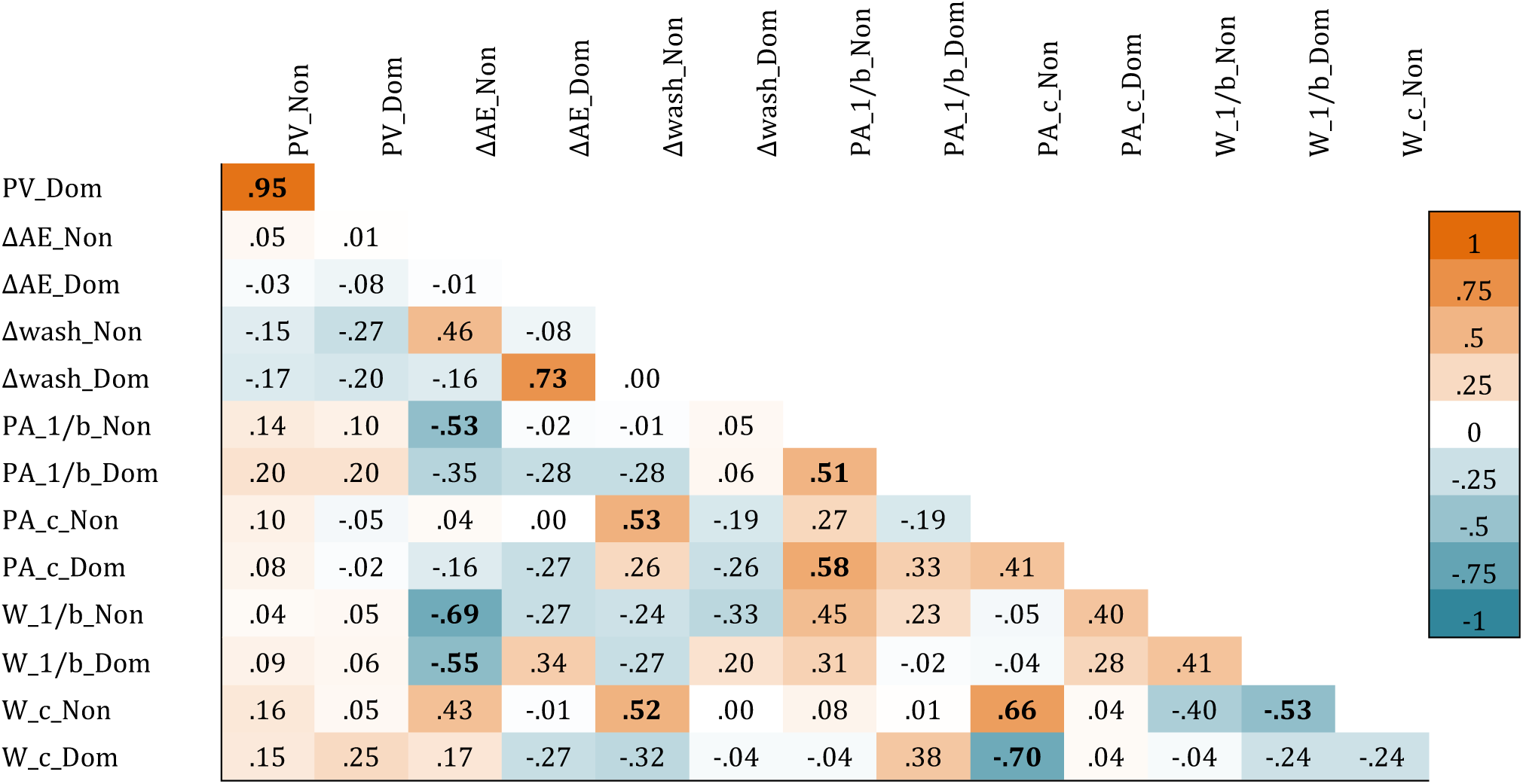
Pearson correlation matrix for controls participants (*n* = 18; B).). Significant correlations (i.e. p < .05) are presented in boldface. ΔAE = endpoint errors from prism adaptation after-effects Open-loop Block, after subtracting baseline pointing error; Δwash = endpoint errors from retention Open-loop Block, after subtracting baseline pointing error; Dom = dominant limb; PV = peak velocity; Non = Non-dominant limb; PA = Prism adaptation; W = Washout.

### 6.3. Supplementary Text T1: Summary statistics

#### 6.3.1. Peak velocity

People with CRPS did not differ from controls in the speed at which they performed pointing movements, although there was a tendency for people with CRPS to move slower when using their affected limb. That is, there was no evidence of a difference between Groups (CRPS *M* = 1533.7 mm/s, *SD* = 343.9; controls *M* = 1575.9 mm/s, *SD* = 467.0), or an effect of Hand (affected/non-dominant *M* = 1545.4 mm/s, *SD* = 413.8; non-affected/dominant *M* = 1563.4 mm/s, *SD* = 412.4) on peak velocity averaged across all open-loop and closed-loop trials, *Fs*(1, 30) ≤ 0.60, *p*s ≥ .443, ƞ^2^ _p_≤ .02. There was, however, a significant interaction between Group and Hand, *F*(1, 30) = 9.05, *p* = .005, ƞ^2^ _p_ = .23. This interaction appeared to be driven by a smaller peak velocity between the affected side (*M* = 1472.6 mm/s, *SD* = 344.8) and the non-affected side (*M* = 1588.0 mm/s, *SD* = 334.0) for people with CRPS, whereas there was less of a difference between peak velocity for the non-dominant (*M* = 1608.2 mm/s, *SD* = 455.9) and dominant (*M* = 1543.2 mm/s, *SD* = 475.9) hands for controls. None of the differences between conditions was significant after correcting for multiple comparisons, *t*s(30) ≤ 2.36, *p*s_adjusted_ ≥ .100, *d*s ≤ 0.84.

#### 6.3.2. Peak acceleration

Peak acceleration was comparable between people with CRPS and controls, although it was lower when people with CRPS used their affected hand. That is, there were no significant differences between Groups (CRPS *M* = 1058.2 mm/s^2^, *SD* = 3995.8; controls *M* = 10968.1 mm/s^2^, *SD* = 6002.6), or between the Hand used (affected/non-dominant *M* = 10437.8 mm/s^2^, *SD* = 5046.2; non-affected/dominant *M* = 11111.7 mm/s^2^, *SD* = 5212.7) on peak acceleration averaged across all trials, *Fs*(1, 30) ≤ 2.55, *p*s ≥ .121, ƞ^2^ _p_≤ .08. There was a significant interaction between Group and Hand, *F*(1, 30) = 5.99, *p* = .021, ƞ^2^ _p_= .17. Follow-up analyses suggested that this interaction was driven by people with CRPS having lower peak acceleration when using the affected Hand (*M* = 9648.1 mm/s^2^, *SD* = 3683.7) and the non-affected Hand (*M* = 11412.9 mm/s^2^, *SD* = 4078.6), *t*(30) = 2.70, *p*_*adjusted*_ = .044, *d* = 0.99. There were no significant difference in peak acceleration between the non-dominant (*M* = 11118.0 mm/s^2^, *SD* = 5891.8) and dominant (*M* = 10816.5 mm/s^2^, *SD* = 6109.6) Hand for controls, and or any differences between Groups that depended on the Hand, *t*s(30) ≤ 1.10, *p*s_adjusted_ ≥ .837, *d*s ≤ 0.40.

#### 6.3.3. Baseline closed-loop pointing errors

Endpoint errors during closed-loop pointing were comparable between people with CRPS and controls. There were no significant main effects of Group (CRPS *M* = −0.01°, *SD* =0.43; controls *M* = −0.05°, *SD* = 0.39), or Hand (affected/non-dominant *M* = −0.12°, *SD* = 0.38; non-affected/dominant *M* = 0.06°, *SD* =0.41), or any interactions on endpoint errors for baseline closed-loop trials, *Fs*(1, 31) ≤ 2.69, *p*s ≥ .111, ƞ^2^_p_ ≤ .08.

### 6.4. Supplementary Text T3: Model fit

#### 6.4.1. Exponential decay of endpoint errors during prism exposure

Before analysing the constants derived from the fitted models, we analysed the model fit. The model failed to converge, or there was no exponential fit for one person with CRPS (non-affected hand), and for one control (dominant hand). Next, we compared the model fit parameters between Groups and Hand, for those cases where the model did converge and there was an exponential decay (CRPS n = 14; controls n = 17). The results suggested that the models were not found to differ across Groups and Hand. That is, the prediction errors (i.e. the root-mean-square error [RMSE]) which indicated the mean distance from a predicted value to an observed value, for individually fitted models was not significantly different between Groups, (CRPS *M*_RMSE_ = 1.20, *SD* = 0.67; controls *M*_RMSE_ = 0.98, *SD* = 0.72), Hand (affected/non-dominant *M*_RMSE_ = 1.09, *SD* = 0.57; non-affected/dominant *M*_RMSE_ = 1.08, *SD* =0.82), and there was no significant interaction between the two variables, *Fs*(1, 29) ≤ 1.70, *p*s ≥ .202, ƞ^2^ _p_ ≤ .06. Similarly, there were no significant differences in how much variance was explained by the models (i.e. the *adj. R*^*2*^) between Groups (CRPS *M*_*adj.R2*_ = .47, *SD* = .26; controls *M*_*adj.R2*_ = .54, *SD* = .26), Hand (affected/non-dominant *M*_*adj.R2*_ = .52, *SD* = .25; non-affected/dominant *M*_*adj.R2*_ = .50, *SD* = .27), and there was no significant interaction between the two variables, *Fs*(1, 29) ≤ 0.52, *p*s ≥ .475, ƞ^2^_p_ ≤ .02. As there was no clear difference in the model fits between people with CRPS and controls, or any clear differences depending on the hand used, we proceeded to analyse the constants derived from the models (i.e. 1/*b*, and *c*; Fig. 4).

#### 6.4.2. Exponential decay of endpoint errors during washout

Prior to analysing the constants from the exponential decay function, we analysed the model fit. We were unable to fit an exponential decay function for two participants with CRPS, both for their affected hand. For those cases where the model did converge and there was an exponential decay (CRPS n = 13; controls n = 18), we compared the model fit parameters between Groups and Hand. The prediction error (i.e. RMSE) did not differ between people with CRPS and controls, or for either hand. That is, there was no significant main effect of Group (CRPS *M*_RMSE_ = 0.77, *SD* = 0.35; controls *M*_RMSE_ = 0.59, *SD* = 0.29), Hand (affected/non-dominant *M*_RMSE_ = 0.64, *SD* = 0.27; non-affected/dominant *M*_RMSE_ = 0.69, *SD* = 0.37), and no significant interactions on RMSE, *Fs*(1, 29) ≤ 2.03, *p*s ≥ .165, ƞ^2^ _p_≤ .07. These results suggest that there was no difference in the prediction error between Groups or the hand used. We then analysed how much variance was explained by the models (i.e. the *adj. R*^2^). There were no significant differences between Groups (CRPS *M*_*adj.R2*_ = .34, *SD* = .23; controls *M*_*adj.R2*_ = .42, *SD* = .30), *F*(1, 29) = 0.43, *p* = .517, ƞ^2^_p_ = .01. There was a tendency for models to explain a greater proportion of the variance for models fitted to data from the non-affected/dominant hand (*M*_*adj.R2*_ = .41, *SD* = .27) than the affected/non-dominant hand (*M*_*adj.R2*_ = .35, *SD* = .28), although not statically significant, *F*(1, 29) = 3.16, *p* = .086, ƞ^2^_p_ = .10. Neither did we find any evidence that this tendency varied between Groups, as there was no significant interaction between Group and Hand on *adj. R*^2^, *Fs*(1, 29) ≤ 1.14, *p*s ≥ .343, ƞ^2^_p_ ≤ .04. Therefore, as there was no clear difference in the model fits between people with CRPS and controls, and the tendency for the models to explain a greater proportion of the variance for the non-affected/dominant hand did not vary between groups, we proceeded to analyse the constants derived from the models (i.e. 1/*b*, and *c*; Fig. 5).

#### 6.4.3. Feedforward motor control

Prior to analysing the detrended data for initial trajectory orientations, we inspected the model fit. We were unable to fit initial trajectory orientations to the exponential decay function for one person with CRPS (non-affected hand), and two controls (one non-dominant hand, one dominant hand). For those cases where we were able to fit an exponential decay to their initial trajectory orientation, we compared the model fit parameters between Groups and Hand. In general, the models fitted to the initial trajectory orientations had greater prediction error (*M*_RMSE_ = 10.58, *SD* = 4.72) and explained less of the variance (*M*_*adj.R2*_ = .02, *SD* = .06) than the models fitted to endpoint errors (*M*_*RMSE*_ = 1.09, *SD* = 0.70; *M*_*adj.R2*_ = .51, *SD* = .26), which indicates that the exponential decay was a better fit for endpoint errors than for initial trajectory orientations. For initial trajectory orientations there was a tendency for the prediction error of the model to be greater for controls (*M*_RMSE_ = 11.80, *SD* = 5.90) than people with CRPS (*M*_RMSE_ = 9.24, *SD* = 2.40), although not significant, *F*(1, 28) = 3.19, *p* = .085, ƞ^2^_p_ = .10. There was no significant difference in RMSE between the Hand used (affected/non-dominant *M*_*RMSE*_ = 10.10, *SD* = 3.42; non-affected/dominant *M*_*RMSE*_ = 11.08, *SD* = 5.83), and there was no significant interaction with Group and Hand, *Fs*(1, 28) ≤ 0.80, *p*s ≥ .379, ƞ^2^ _p_ ≤ .03. This suggests that the prediction error did not vary depending on the hand used, although there was a trend for models to make greater prediction errors for controls participants than people with CRPS. Next we analysed how much of the variance in the data was accounted for by the models (i.e. *adj. R*^2^). There were no significant differences between Groups (CRPS *M*_*adj.R2*_ = .04, *SD* = .08; controls *M*_*adj.R2*_ = .01, *SD* = .04), or Hand (affected/non-dominant *M*_*adj.R2*_ = .03, *SD* = .07; non-affected/dominant *M*_adj.R2_ = .02, *SD* = .06) on *adj. R*^*2*^, and no significant interaction, *Fs*(1, 28) ≤ 2.39, *p*s ≥ .134, ƞ^2^ _p_ ≤ .08. This suggests that the amount of variance explained by the models did not differ between people with CRPS and controls, or depending on the Hand used. However, it should be noted that the amount of variance explained by an exponential fit for initial trajectory orientations were substantially lower than that of endpoint errors.

### 6.5. Supplementary Text T3: Exploratory analysis – Correlational analyses

For controls, the difference in open-loop endpoint errors during prism exposure compared to baseline showed a strong, significant correlation with the difference in open-loop endpoint errors during washout compared to baseline for the dominant hand, *r* = .73, *p* < .001, but this was not observed for the non-dominant hand, *r* = .46, *p* = .056. For people with CRPS, the difference in open-loop endpoint errors during prism exposure compared to baseline showed a strong, significant correlation with the difference in open-loop endpoint errors during washout compared to baseline for the affected hand, *r* = .82, *p* = .002, but not for the non-affected hand, *r* = .36, *p* = .243.

For people with CRPS, the severity of their conditions showed moderate to strong correlations with upper limb disability, neuropathic type pain, body representation disturbance, and “neglect-like symptoms”, *r*s ≥ .49, *p*s ≤ .047. The latter was also negatively correlated with the duration of CRPS. Furthermore, baseline pain was associated with fear of movement, upper limb disability, body representation disturbance, and “neglect-like symptoms”.

### 6.6. Supplementary Text T4: Exploratory analysis – Covariate analyses

People with unilateral CRPS have been reported to have bilateral proprioceptive deficits (Bank et al., 2013). In previous research, absolute pointing errors made with the unseen hand(s) have been interpreted as evidence of deficits in arm position sense in people with CRPS (Lewis et al., 2010). Therefore, we used the absolute endpoint error during the baseline Open-loop Block (“Absolute Baseline Error”) as a measure of proprioceptive accuracy. Our results, however suggest that there were no differences in proprioceptive accuracy between people with CRPS and controls, *Fs*(1, 28) ≤ 0.06, *p*s ≥ .805, ƞ^2^_p_ < .01. When we re-ran our primary analysis (3.3.1.) of open-loop endpoint errors using ANCOVAs with proprioceptive accuracy included as a covariate, we found that proprioceptive accuracy did not influence our results, *Fs*(2, 62) ≤ 1.03, *p*s ≥ .362, ƞ^2^ _p_≤ .03. Next, because more of our participants with CRPS than controls were exposed to leftward-shifting prisms, we considered that the direction of the prismatic shift might have influenced the degree of sensorimotor realignment, as a difference between adapting to leftward and rightward shifting lenses has been reported previously (Redding & Wallace, 2009). We also did not observe any influence of the direction of the prismatic shift on our results from follow-up ANCOVAs, *Fs*(2, 62) ≤ 0.62, *p*s ≥ .544, ƞ^2^ _p_≤ .02, which suggests that our findings are unlikely to be due to a greater number of people with CRPS being exposed to leftward shifting prisms than controls. Therefore, it does not seem likely that our findings can be attributed to differences in proprioceptive abilities, the direction of the prismatic shift, and/or the counterbalancing order.

We considered that the counterbalancing order might influence our results, as inter-limb transfer has previously been found to be greater when the dominant hand is adapted first, compared to the non-dominant hand (Redding & Wallace, 2008, 2011). When we reanalysed the endpoint errors with Counterbalancing Order (affected/non-dominant first, non-affected/dominant first), Hand, and Open-loop Block as independent variables. There was no significant main effect of Counterbalancing Order on endpoint errors, and/or no significant interactions with Hand, or Open-loop Block, *Fs*(2, 62) ≤ 2.64, *p*s ≥ .080, ƞ^2^ _p_≤ .08. These results therefore suggest that there was no significant influence of inter-limb transfer on endpoint errors.

#### 6.7. Supplementary Text T5: Exploratory analysis – Kinematic changes

As we anticipated the kinematic data to be noisy, we first analysed the strength of individual correlations for each Hand, pooling the data for each Group. That is, we compared *t*-values to zero for each Hand, averaged across all participants. The t values for participants’ non-affected/dominant hand significantly deviated from 0, indicating the presence of a linear relationship between endpoint errors (trial_n_) and the subsequent change in initial trajectory orientation (trial_n+1_) (*M*_t_ = −0.46, *SD* = 1.23), *t*(32) = 2.15, *p* = .039, *d* = 0.37. This association suggests that when participants made endpoint errors in a given direction they adjusted their movement plan in the opposite direction on the subsequent trial (e.g. if an endpoint error was made towards the right, the subsequent movement was angled more towards the left). We did not observe evidence of such a linear relationship when people used their affected/non-dominant hand (*M*_*t*_ = −0.18, *SD* = 1.24), *t*(31) = 0.83, *p* = .416, *d* = 0.15. Therefore, the data pooled across all participants provides evidence that feedforward motor control was used to reduce trial-by-trial endpoint errors relative to the previous trial for the non-affected/dominant hand - but not the affected/non-dominant hand - during early prism exposure trials.

Unilateral Kolmogorov-Smirnov distribution tests (Vindras et al., 2012) on the *p*-values obtained from individual correlations analyzed pooled across Groups, reveled no evidence that *p*-values were biased towards zero for either hand, *D*^+^ ≤ 0.18, *ps* ≥ .104. These findings suggest that the *p*-values for the correlation between endpoint errors for early trials (*n*) and the change in initial trajectory orientation on the next trial (i.e. trial_*n*+1_ - trial_*n*_), were not biased towards zero.

Taken together, there was evidence that feedforward motor control was used to reduce endpoint errors during early prism exposure when participants used the non-affected/dominant hands, but only from the analysis of *t*-values. There was no evidence of such use of feedforward motor control from *t*-values when participants used the affected/non-dominant hands. Therefore, the evidence of feedforward motor control during early trials for the non-affected/dominant hands should be interpreted with caution.

